# A Computational Model of Oxidative Stress in a Human Ventricular Myocyte

**DOI:** 10.64898/2026.09.17.752441

**Authors:** Hannah M. Zukowski, Gonzalo Hernandez Hernandez, Pei Chi Yang, Colleen E. Clancy

## Abstract

Mitochondrial reactive oxygen species (ROS) are implicated in cardiac dysfunction, but complex dynamic interactions between ROS, intracellular calcium, and electrophysiology make it difficult to resolve mechanisms experimentally. We developed the Zukowski excitation-contraction-mitochondrial-ROS (ECM-ROS) model, a computational model that couples human ventricular electrophysiology with mitochondrial calcium handling, energetics, ROS production, and ROS dependent modulation of the ryanodine receptor, SERCA, L-type calcium current, and late sodium current. Model predictions were evaluated against independent experimental measurements of ROS induced changes in action potential duration and intracellular calcium. We then used the model to characterize ROS calcium feedback across physiological pacing rates and applied it to doxorubicin induced oxidative stress. Increasing cytosolic calcium produced nonlinear amplification of mitochondrial activity and cytosolic ROS, with greater amplification at faster pacing rates. Moderate ROS elevation initially enhanced calcium release, whereas greater oxidative stress depleted sarcoplasmic reticulum calcium stores. During simulated doxorubicin exposure, faster pacing lowered the level of oxidative stress required to produce calcium dysregulation and action potential prolongation. Mitochondrial ROS and calcium accumulation preceded detectable electrophysiological remodeling. In populations incorporating variability in ion channel expression, neither doxorubicin induced oxidative stress nor subtle pharmacological I_Kr_ block alone produced early afterdepolarizations. However, early afterdepolarizations were predicted to emerge with combined perturbations. These findings establish a computational framework for investigating bidirectional coupling among mitochondrial function, ROS, calcium handling, and human cardiac electrophysiology. The model generates testable predictions regarding the early detection of oxidative cardiac injury and the physiological conditions that increase susceptibility to arrhythmia.

**First author profile:** Hannah Zukowski is a PhD candidate in Biomedical Engineering at the University of California, Davis, where she works with Dr. Colleen E. Clancy in computational cardiovascular physiology. She earned her B.S. in Mechanical Engineering from Trinity College, where her early work in computational modeling led to an interest in applying engineering approaches to biological systems. Her doctoral research focuses on mechanistic models of mitochondrial bioenergetics, reactive oxygen species, and cardiac electrophysiology to investigate oxidative stress-induced cardiotoxicity. Her future research interests include multiscale and patient-specific cardiovascular modeling to improve the prediction of cardiac dysfunction and treatment response.

**KEY POINTS SUMMARY:**

- Oxidative stress can disrupt calcium regulation and electrical activity in human heart cells, but the interactions among these processes are difficult to study experimentally.
- We developed a computational model that connects mitochondrial function and reactive oxygen species with calcium regulation and electrical activity in human heart cells.
- The model predicts that increases in calcium can amplify mitochondrial activity and reactive oxygen species, particularly at faster heart rates.
- During simulated exposure to the chemotherapy drug doxorubicin, reactive oxygen species accumulation developed before detectable changes in electrical activity. Faster heart rates also increased the effects of oxidative stress.
- The model provides a framework to investigate how oxidative stress contributes to cardiac dysfunction and to identify conditions that may increase susceptibility to abnormal electrical activity.

## INTRODUCTION

Oxidative stress is widely implicated in cardiac disease and contributes to arrhythmia, contractile dysfunction, and heart failure across a broad range of conditions including ischemia-reperfusion injury, atrial fibrillation, diabetic cardiomyopathy, and chemotherapy-induced cardiotoxicity (Conklin, 2004; Tsutsui *et al*., 2011; Liu *et al*., 2014; Xie *et al*., 2015; Liguori *et al*., 2018; Peoples *et al*., 2019; D’Oria *et al*., 2020; Rocca *et al*., 2020; Wu *et al*., 2020; Carrasco *et al*., 2021; Fabiani *et al*., 2021; De Geest & Mishra, 2022; Jurcau & Ardelean, 2022; Pfenniger *et al*., 2024). Reactive oxygen species (ROS), produced in part by mitochondria, are coupled to cardiac excitation–contraction coupling (Murphy, 2009; Köhler *et al*., 2014). During each cardiac cycle, myocardial cytosolic calcium enters the mitochondria and stimulates ATP production. Mitochondrial ROS are produced as byproducts of electron transport chain activity. ROS then enter the cytosol and can modulate calcium-handling proteins and ion channels, including the ryanodine receptor (RyR), SERCA, L-type calcium, and late sodium channels (Brookes *et al*., 2004; Zima & Blatter, 2006). Despite the broad recognition that oxidative stress contributes to cardiac pathology, the specific mechanisms by which mitochondrial ROS production translates into altered electrophysiology and arrhythmia risk remain elusive. Progress has been limited, in part, because mitochondrial dysfunction and electrophysiological remodeling are often investigated as separate processes rather than as components of an integrated, dynamically coupled system (Chen & Zweier, 2014; Kohlhaas *et al*., 2017; Popoiu *et al*., 2023). Complex systems such as cardiac ROS production and its downstream effects are ideally suited to computational modeling, which can reveal and test plausible mechanisms, characterize their interactions, and overcome key experimental limitations.

The coupling between mitochondrial ROS production, mitochondrial calcium, and electrophysiology is difficult to study experimentally. Simultaneous, real-time measurement of mitochondrial ROS production, mitochondrial calcium, and action potential morphology within the same beating myocyte is confounded by technical limitations. For example, current ROS measurement indicators lack the species specificity and temporal resolution needed to isolate cause from effect. Moreover, the coupling between ROS generation and downstream effects is essentially a feedback loop that operates on the same beat-to-beat timescale as excitation-contraction coupling (Kalyanaraman *et al*., 2012; Griendling *et al*., 2016). Most experimental studies of ROS effects in the myocardium apply a fixed concentration of an oxidant and report a single endpoint measurement (Boraso & Williams, 1994; Ward & Giles, 1997; Song *et al*., 2006; Xie *et al*., 2009; Fei *et al*., 2017), an approach that cannot capture how electrophysiological and mitochondrial dysfunction evolve as oxidative stress develops.

Computational modeling offers an alternative approach to overcome experimental limitations. Electrophysiology, mitochondrial energetics, and ROS dynamics can be integrated within a single framework to isolate the individual and combined contributions of each process to myocardial phenotype. Simulations can predict mechanisms that link mitochondrial dysfunction to electrophysiological remodeling and test the effects of pacing rate and pharmacological perturbations.

Early computational models established the link between mitochondrial calcium and cardiac energy production. Cortassa et al. (2003) developed a model of mitochondrial calcium handling, the tricarboxylic acid (TCA) cycle, and oxidative phosphorylation that linked cytosolic calcium to mitochondrial ATP production. The model predicted that mitochondrial calcium contributes to the adaptation of energy supply during changes in cardiac workload (Cortassa *et al*., 2003). Cortassa et al. (2004) extended this framework to include mitochondrial ROS production and scavenging. This model reproduced oscillations in mitochondrial membrane potential, NADH, and ROS and demonstrated how ROS balance can regulate mitochondrial function (Cortassa *et al*., 2004). Cortassa et al. (2006) subsequently integrated mitochondrial energetics with a model of ventricular electrophysiology, calcium handling, and contraction to form the excitation-contraction-mitochondrial energetics (ECME) model. ATP, calcium, and sodium linked cellular activity to mitochondrial metabolism. The model reproduced changes in oxygen consumption, mitochondrial NADH, and calcium in response to changes in cardiac workload and provided a framework to study the reciprocal relationship between energy demand and mitochondrial energy production (Cortassa *et al*., 2006). Subsequent analysis of the ECME model demonstrated that control of mitochondrial respiration is distributed across mitochondrial and cytoplasmic processes and shifts with cardiac workload (Cortassa *et al*., 2009). Zhou et al. (2009) extended the ECME model to include mitochondrial ROS-induced ROS release and a sarcolemmal ATP-sensitive potassium current (I_KATP_). The model linked mitochondrial oxidative stress to membrane electrophysiology through changes in cellular energy state. Simulations predicted that ROS-induced mitochondrial depolarization reduced the cytosolic ATP/ADP ratio, activated I_KATP_, and shortened action potential duration (Zhou *et al*., 2009). Li et al. (2015) further extended the ECME-RIRR framework to include local calcium control, spatial ROS transport between mitochondria and the sarcoplasmic reticulum, and direct ROS-dependent regulation of RyR and SERCA. The model predicted that mitochondrial ROS bursts increase cytosolic calcium through RyR activation and SERCA inhibition, which can produce abnormal action potentials. The timing and magnitude of ROS release and the distance between mitochondria and the sarcoplasmic reticulum determined the resulting electrical phenotype (Li *et al*., 2015).

Other models have focused on different levels of mitochondrial detail. Gauthier et al. (2013) developed a detailed model of ROS production from complexes I and III of the electron transport chain and mitochondrial ROS scavenging (Gauthier *et al*., 2013). In contrast, Senneff and Lowery (2022) used a reduced representation of mitochondrial energetics to couple mitochondrial calcium and ATP production to excitation-contraction coupling in skeletal muscle (Senneff & Lowery, 2022). Together, these models established important links between mitochondrial energetics, ROS, calcium handling, and cardiac electrophysiology. However, no existing model integrates these processes within a human ventricular myocyte. The most comprehensive integrated cardiac models were developed for guinea pig ventricular myocytes and require detailed descriptions of mitochondrial metabolism and ROS dynamics. This model complexity limits their use for large-scale population simulations. A reduced human ventricular model that retains the key feedback mechanisms between mitochondrial ROS, calcium handling, and electrophysiology remains needed.

Here, we address this gap with the Zukowski excitation-contraction-mitochondrial-ROS (ECM-ROS) model, which couples the O’Hara-Rudy (ORd) human ventricular electrophysiological model (O’Hara *et al*., 2011) to reduced descriptions of mitochondrial energetics and ROS dynamics. The mitochondrial module describes mitochondrial membrane potential, calcium handling, and electron transport chain activity through a set of mitochondrial fluxes. Mitochondrial ROS production and scavenging are coupled to this energetic state. Cytosolic ROS directly modulate calcium-handling proteins and ion channels consistently implicated in oxidative injury which include the RyR, SERCA, and the L-type calcium (I_CaL_) and late sodium currents (I_NaL_). The model design retains the essential bidirectional coupling between electrophysiology, mitochondrial calcium, and ROS and remains computationally efficient to support large-scale population simulations of arrhythmia susceptibility.

We applied the Zukowski ECM-ROS model to doxorubicin (DOX) as a clinically relevant test case to investigate oxidative stress-induced arrhythmia susceptibility (Volkova & Russell, 2012; Huang *et al*., 2022; Kong *et al*., 2022; Bhutani *et al*., 2025). DOX is a widely used chemotherapeutic agent whose clinical utility is limited by dose-dependent cardiotoxicity, with greater cumulative exposure associated with greater mitochondrial dysfunction (Dudka *et al*., 2012; Cappetta *et al*., 2017; Li *et al*., 2025). One mechanism by which DOX generates ROS occurs at Complex I of the electron transport chain. DOX undergoes one-electron reduction and forms a semiquinone radical that reacts with molecular oxygen to produce superoxide while regenerating DOX. This redox cycling process increases mitochondrial superoxide generation in a dose-dependent manner (Davies & Doroshow, 1986; Doroshow & Davies, 1986). DOX further impairs antioxidant defense, reducing the scavenging capacity available to counteract excess superoxide production (Doroshow *et al*., 1980; Li & Singal, 2000). The combination of enhanced mitochondrial ROS production and impaired scavenging makes acute DOX cardiotoxicity a direct physiological test of the ROS-calcium feedback loop that the Zukowski ECM-ROS model is designed to capture.

Several questions related to DOX-induced cardiotoxicity remain unresolved from existing experimental and computational studies. First, whether the severity of DOX-induced mitochondrial and electrophysiological dysfunction depends on heart rate is unknown, since most experimental studies are conducted at a fixed pacing rate (Wang & Korth, 1995; George *et al*., 2026). Second, the temporal relationship between mitochondrial dysfunction and electrophysiological remodeling during the development of DOX-induced oxidative stress has not been established, despite its relevance to identification of early disease biomarkers. Current clinical monitoring for DOX cardiotoxicity relies on downstream structural and functional readouts such as reduced ejection fraction (Plana *et al*., 2014) and QT prolongation (Yeh *et al*., 2004; Octavia *et al*., 2012). Third, whether DOX-induced oxidative stress alone is sufficient to trigger arrhythmia, or requires an additional stressor, remains untested. This question is clinically relevant given that cancer patients treated with DOX frequently receive medications that block the rapid delayed rectifier potassium current and prolong the QT interval (Agnihotri *et al*., 2024; Ramasubbu *et al*., 2025).

The Zukowski ECM-ROS model represents a first-stage integrated model of human ventricular electrophysiology, mitochondrial energetics, and ROS dynamics. We developed the model by coupling ORd to a modified mitochondrial formulation from Senneff and Lowery (2022) and ROS-induced ROS release mechanisms adapted from Cortassa et al. (2004). We added ROS-dependent regulation of RyR, SERCA, I_CaL_, and I_NaL_ to couple mitochondrial oxidative stress directly to calcium handling and membrane electrophysiology. This reduced framework provides a human ventricular model of the ROS-calcium feedback loop suitable for population-scale simulations. We used the model to predict the effects of DOX-induced oxidative stress across physiologically relevant pacing rates and to define the temporal relationship between mitochondrial ROS, calcium dysregulation, and electrophysiological remodeling. Finally, we incorporated inter-individual variability in ion channel conductance to test whether DOX-induced oxidative stress, alone or in combination with pharmacological I_Kr_ block, is sufficient to produce arrhythmogenic afterdepolarizations.

## METHODS

### Computational Model Overview

The Zukowski ECM-ROS model is an integrated computational model of excitation-contraction coupling (EC), mitochondrial (M) energetics, and reactive oxygen species (ROS) dynamics in the human ventricular myocyte. The electrophysiological component is the O’Hara-Rudy dynamic (ORd) model of the undiseased human ventricular myocyte (O’Hara *et al*., 2011). The mitochondrial energetics and ROS components are adapted from the Senneff-Lowery and Cortassa models (Cortassa *et al*., 2004; Senneff & Lowery, 2022). Cytosolic calcium ([Ca^2+^]_i_) couples whole-cell electrophysiology to mitochondrial metabolism and enables beat-to-beat feedback between calcium dynamics, mitochondrial energetics, and ROS production. ROS generated by the mitochondria modulates key ion channels and calcium handling proteins (I_CaL_, I_NaL_, RyR, and SERCA) which creates a bidirectional feedback loop between mitochondrial dysfunction and electrophysiological remodeling. A full description of mitochondria and ROS model parameters is provided in **Table 1**.

**Table 1:** Mitochondrial module parameters of the Zukowski ECM-ROS model. Parameters are grouped by functional subsystem. ORd electrophysiology parameters are published in O’Hara et al. (2011) and are not listed here.

| Symbol | Description | Value | Units | Source |
| --- | --- | --- | --- | --- |
| *General Mitochondrial Parameters* |  |  |  |  |
| $C_p$ | Mitochondrial membrane capacitance (divided by F) | $1.8 \times 10^{-3}$ | mM/mV | Senneff & Lowery, 2022 |
| $a_1$ | Scaling factor between NADH consumption and $\Delta\psi$ | 120.0 | Dimensionless | Senneff & Lowery, 2022 |
| $a_2$ | Scaling factor between ATP production and $\Delta\psi$ | 3.43 | Dimensionless | Senneff & Lowery, 2022 |
| $\partial$ | Fraction of free (unbuffered) mitochondrial $Ca^{2+}$ | $3 \times 10^{-4}$ | Dimensionless | Cortassa et al., 2003 |
| $[NAD]_{total}$ | Total mitochondrial pyridine nucleotide (NAD) | 2.970 | mM | Senneff & Lowery, 2022 |
| *Mitochondrial Permeability Transition Pore ($J_{mPTP}$)* |  |  |  |  |
| $p_3$ | Voltage dependence coefficient of mPTP | 0.075 | mV<sup>-1</sup> | Senneff & Lowery, 2022 |
| $k_{mPTP}$ | Rate constant of Ca<sup>2+</sup> leak (mPTP) | 8.0 x 10<sup>-6</sup> | mM/ms | TUNED from Senneff & Lowery, 2022 |
| *Mitochondrial Ca<sup>2+</sup> Uniporter (J<sub>MCU</sub>)* |  |  |  |  |
| $V_{MCU}$ | Maximum uniporter Ca<sup>2+</sup> transport rate | 0.0275 | mM/ms | Cortassa et al., 2003 |
| $K_{trans}$ | Dissociation constant (K<sub>d</sub>) for translocated Ca<sup>2+</sup> | 0.019 | mM | Cortassa et al., 2003 |
| $K_{act}$ | Activation constant for uniporter | 3.8 x 10<sup>-4</sup> | mM | Cortassa et al., 2003 |
| $L$ | Equilibrium constant (K<sub>eq</sub>) for uniporter conformational transitions | 110 | Dimensionless | Cortassa et al., 2003 |
| $n_a$ | Uniporter activation cooperativity | 2.8 | Dimensionless | Cortassa et al., 2003 |
| *Mitochondrial Na<sup>+</sup>/Ca<sup>2+</sup> Exchanger (J<sub>mNCX</sub>)* |  |  |  |  |
| $V_{NCX}$ | Maximum mitochondrial NCX flux | 1.0 x 10<sup>-4</sup> | mM/ms | Cortassa et al., 2003 |
| $\Delta\psi^0$ | Membrane potential offset | 91 | mV | Cortassa et al., 2003 |
| $K_{Na}$ | Antiporter Na<sup>+</sup> constant | 9.4 | mM | Cortassa et al., 2003 |
| $K_{Ca}$ | Antiporter Ca<sup>2+</sup> constant | 3.75 x 10<sup>-4</sup> | mM | Cortassa et al., 2003 |

| *Mitochondrial Metabolism ($J_{PDH}$, $J_{AGC}$)* |  |  |  |  |
| --- | --- | --- | --- | --- |
| $K_{AGC}$ | Ca<sup>2+</sup> dissociation constant for AGC | $0.14 \times 10^{-3}$ | mM | Senneff & Lowery, 2022 |
| $p_4$ | Voltage dependence coefficient of AGC activity | 0.01 | mV<sup>-1</sup> | Senneff & Lowery, 2022 |
| $q_1$ | Michaelis-Menten-like constant for NAD<sup>+</sup> consumption by Krebs cycle | 0.2244 | dimensionless | Cortassa et al., 2003 |
| $q_2$ | S<sub>0.5</sub> for Ca<sup>2+</sup> activation of Krebs cycle | $0.1 \times 10^{-3}$ | mM | Senneff & Lowery, 2022 |
| $V_{AGC}$ | Rate constant of AGC (NADH production via malate-aspartate shuttle) | $0.025 \times 10^{-3}$ | mM/ms | Senneff & Lowery, 2022 |
| $k_{GLY}$ | Empirical velocity of glycolysis | $4.5 \times 10^{-4}$ | mM/ms | Wacquier et al., 2016 |
| *Oxidative Phosphorylation ($J_{ETC}$, $J_{F1F0}$, $J_{ANT}$, $J_{HLeak}$)* |  |  |  |  |
| $q_3$ | Michaelis-Menten constant for NADH consumption by ETC | $100 \times 10^{-3}$ | mM | Senneff & Lowery, 2022 |
| $q_4$ | Voltage dependence coefficient 1 of ETC activity | 177.0 | mV | Senneff & Lowery, 2022 |
| $q_5$ | Voltage dependence coefficient 2 of ETC activity | 5.0 | mV | Senneff & Lowery, 2022 |
| $q_6$ | ATP inhibition constant of F1F0 | $10000 \times 10^{-3}$ | mM | Senneff & Lowery, 2022 |
| $q_7$ | Voltage dependence coefficient 1 of F1F0 | 190.0 | mV | Senneff & Lowery, 2022 |
| $q_8$ | Voltage dependence coefficient 2 of F1F0 | 8.5 | mV | Senneff & Lowery, 2022 |
| $q_9$ | Voltage-dependent proton leak coefficient | $0.0020 \times 10^{-3}$ | mM/(ms·mV) | Senneff & Lowery, 2022 |
| $q_{10}$ | Voltage-independent proton leak rate constant | $-0.030 \times 10^{-3}$ | mM/ms | Senneff & Lowery, 2022 |
| $V_{ANT}$ | Rate constant of adenine nucleotide translocator (ANT) | $8.123 \times 10^{-3}$ | mM/ms | Senneff & Lowery, 2022 |
| $V_{F1F0}$ | Rate constant of F1F0-ATPase | $3.6 \times 10^{-3}$ | mM/ms | Senneff & Lowery, 2022 |
| $k_{m,ADP}$ | Michaelis-Menten constant for ANT | $3.5 \times 10^{-3}$ | mM | Senneff & Lowery, 2022 |
| $\theta$ | ANT voltage distribution parameter | 0.35 | dimensionless | Senneff & Lowery, 2022 |
| $V_{ETC}$ | Rate constant of NADH oxidation by ETC | $0.764 \times 10^{-3}$ | mM/ms | Senneff & Lowery, 2022 |
| **ROS Generation and Transport (IMAC)** |  |  |  |  |
| $ETCROSLeak$ | Fraction of $J_{ETC}$ diverted to mitochondrial $O_2^{\cdot -}$ | 0.40–0.60 | dimensionless | Varied in simulations |
| $G_L$ | Leak conductance for IMAC | $7.82 \times 10^{-8}$ | mM/(ms·mV) | Cortassa et al., 2004 |
| $G_{\max}$ | Integral conductance of IMAC at saturation | $7.82 \times 10^{-6}$ | mM/(ms·mV) | Cortassa et al., 2004 |
| $\kappa$ | Steepness factor for IMAC voltage dependence | $70 \times 10^{-3}$ | mV<sup>-1</sup> | Cortassa et al., 2004 |
| $E_{m,ROS}$ | Membrane potential at half-saturation of IMAC | 4.0 | mV | Cortassa et al., 2004 |
| $b_{ROS}$ | Activation factor for cytosolic O<sub>2</sub><sup>-</sup> on IMAC | $1.0 \times 10^4$ | dimensionless | Cortassa et al., 2004 |
| $K_{cc}$ | Activation constant of IMAC by cytosolic O<sub>2</sub><sup>-</sup> | 0.01 | mM | Cortassa et al., 2004 |
| $j$ | Fraction of IMAC conductance for O<sub>2</sub><sup>-</sup> transport | 0.1 | dimensionless | Cortassa et al., 2004 |
| **ROS Scavenging (SOD, CAT, GPx, GR)** |  |  |  |  |
| $k_{SOD}^1$ | Second-order rate constant: oxidized SOD + O<sub>2</sub><sup>-</sup> | $1.2 \times 10^3$ | mM<sup>-1</sup>ms<sup>-1</sup> | (Zhou et al., 2009 |
| $k_{SOD}^3$ | Second-order rate constant: reduced SOD + O<sub>2</sub><sup>-</sup> | 24 | mM<sup>-1</sup>ms<sup>-1</sup> | Zhou et al., 2009 |
| $k_{SOD}^5$ | First-order rate constant: inactive → active oxidized SOD | $0.25 \times 10^{-3}$ | ms<sup>-1</sup> | Zhou et al., 2009 |
| $E_{SOD}^T$ | Intracellular SOD concentration | $1.5 \times 10^{-3}$ | mM | Cortassa et al., 2004 |
| $K_i^{H_2O_2}$ | H<sub>2</sub>O<sub>2</sub> inhibition constant for SOD | 0.5 | mM | Zhou et al., 2009 |
| $k_{CAT}^1$ | Rate constant of catalase | 17.0 | mM<sup>-1</sup>ms<sup>-1</sup> | Zhou et al., 2009 |
| $E_{CAT}^T$ | Intracellular catalase concentration | 0.01 | mM | Zhou et al., 2009 |
| $fr$ | H<sub>2</sub>O<sub>2</sub> inhibition factor for catalase | 0.05 | dimensionless | Zhou et al., 2009 |
| $E_{GPX}^T$ | Intracellular GPx concentration | 0.01 | mM | Zhou et al., 2009 |
| $\Phi_1$ | GPx activity constant | $0.5 \times 10^{-2}$ | mM·ms | Zhou et al., 2009 |
| $\Phi_2$ | GPx activity constant | 0.75 | mM·ms | Zhou et al., 2009 |
| $k_{GR}^1$ | Rate constant of glutathione reductase | $5.0 \times 10^{-3}$ | ms<sup>-1</sup> | Zhou et al., 2009 |
| $E_{GR}^T$ | Intracellular GR concentration | 0.01 | mM | Zhou et al., 2009 |
| $K_M^{GSSG}$ | GSSG Michaelis constant for GR | 0.06 | mM | Zhou et al., 2009 |
| $K_M^{NADPH}$ | NADPH Michaelis constant for GR | 0.015 | mM | Zhou et al., 2009 |
| $G_T$ | Total glutathione concentration | 1.0 | mM | Zhou et al., 2009 |
| $NADPH$ | Cytosolic NADPH concentration (fixed) | 1.0 | mM | Zhou et al., 2009 |

### Electrophysiology: O’Hara-Rudy Model

The electrophysiological component of the Zukowski ECM-ROS model is the O’Hara-Rudy dynamic (ORd) model of the undiseased human ventricular myocyte (O’Hara *et al*., 2011). The ORd model comprises ordinary differential equations (ODEs) that govern transmembrane voltage, ionic currents (including fast and late sodium current, I_Na_ and I_NaL_; L-type calcium current, I_CaL_; and potassium currents I_Kr_, I_Ks_, and I_K1_), and intracellular ion concentrations in the cytosol, subspace (SS), junctional SR (JSR), and network SR (NSR) compartments. Sarcoplasmic reticulum (SR) calcium cycling is governed by ryanodine receptor release flux (J_RyR_) and SERCA uptake flux (J_up_). The ORd model was implemented as published (O’Hara *et al*., 2011), with one modification to the cytosolic calcium ODE to incorporate mitochondrial calcium fluxes (described in **Model Coupling** section).

### Mitochondrial Energetics Module

The mitochondrial energetics module was adapted from the Senneff-Lowery model of mitochondrial calcium handling and oxidative phosphorylation in skeletal muscle (Senneff & Lowery, 2022), which draws flux formulations from Wacquier et al. (2016) and the Cortassa et al. (2003) integrated model of cardiac mitochondrial energy metabolism and calcium dynamics (Cortassa *et al*., 2003; Wacquier *et al*., 2016). The Senneff-Lowery model was selected as the foundation for the mitochondrial module because it provides a simplified, yet physiologically grounded representation of calcium-regulated mitochondrial energetics, sufficient to capture beat-to-beat coupling between calcium dynamics, oxidative phosphorylation, and ROS generation without the full complexity of the TCA cycle intermediate dynamics described in Cortassa et al. (2003). The original Senneff-Lowery model describes mitochondrial calcium handling and associated energetic fluxes across two spatial sub compartments, a terminal space proximal to the SR calcium release site and a bulk space distal from the release site (Senneff & Lowery, 2022). In the Zukowski ECM-ROS model, all mitochondrial flux was computed in a single bulk compartment that represents mean mitochondrial matrix conditions.

### Mitochondrial Calcium Handling

Mitochondrial calcium ([Ca^2+^]_m_) dynamics are governed by three fluxes: uptake through the mitochondrial calcium uniporter (J_MCU_), extrusion through the mitochondrial Na^+^/Ca^2+^ exchanger (J_NCX_), and flux through the mitochondrial permeability transition pore (J_mPTP_) (Senneff & Lowery, 2022). The ODE governing mitochondrial calcium is:

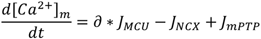

Mitochondrial calcium concentration was scaled by ∂, which represents the fraction of free mitochondrial calcium following mitochondrial calcium buffering reactions.

Mitochondrial calcium uptake through the uniporter (J_MCU_) is driven by the mitochondrial membrane potential (ΔΨ) and the cytosolic-to-mitochondrial calcium concentration gradient (Cortassa *et al*., 2003):

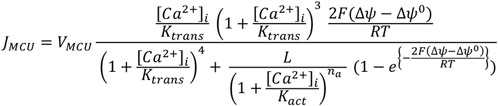

V_MCU_ is the maximal calcium uptake rate, K_trans_ is K_d_ for translocated calcium, K_act_ is an activation constant, and L is the K_eq_ for conformational transitions, n_a_ is the activation cooperativity, and ΔΨ^0^ is the offset membrane potential.

Mitochondrial calcium extrusion through the Na^+^/Ca^2+^ exchanger (J_NCX_) is drive by ΔΨ and the logarithmic ratio of cytosolic to mitochondrial calcium (Cortassa *et al*., 2003) :

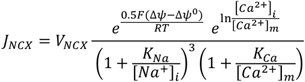

V_NCX_ is the maximal NCX rate, K_Na_ is the antiporter sodium constant, K_Ca_ is the antiporter calcium constant, and ΔΨ^0^ is the offset membrane potential.

Flux through the mitochondrial permeability transition pore (J_mPTP_) is driven by the cytosolic-to-mitochondrial calcium gradient and is modulated by ΔΨ (Senneff & Lowery, 2022):

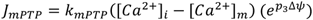

K_mPTP_ is the rate constant of bidirectional calcium leak from the mitochondria and p_3_ is the voltage dependence coefficient of calcium leak.

### Metabolic Fluxes, NADH Production, and Oxidative Phosphorylation

Activation of the electron transport chain (ETC) requires a pool of mitochondrial NADH. NADH accumulation is initiated through activation of the aspartate-glutamate carrier (J_AGC_). J_AGC_ is activated by cytosolic calcium, with maximal flux rate V_AGC_, calcium dissociation constant K_AGC_, and voltage dependence with coefficient p_4_ (Senneff & Lowery, 2022):

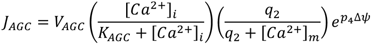

Following AGC activation, the pyruvate dehydrogenase complex (J_PDH_) initiates the reduction of NAD^+^ to NADH as part of the Krebs cycle (Senneff & Lowery, 2022). Mitochondrial NAD^+^ is modeled by a conservation equation where <NAD>_total_ is the total concentration of oxidized and reduced NAD ions within the mitochondria set to 2.970mM (Senneff & Lowery, 2022):

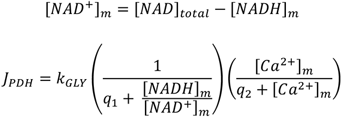

J_PDH_ captures both NAD^+^ reduction scaled by glycolytic rate k_GLY_, with q_1_ as a Michaelis-like constant for NAD^+^ consumption and mitochondrial calcium-dependent activation of the Krebs cycle, where q_2_ is the half-activation constant. Notably, AGC and PDH activity share a calcium-dependent mechanism through q_2_.

The time rate change of mitochondrial NADH (NADH_m_) is governed by the balance of NADH production through J_PDH_, J_AGC_, and NADH consumption by ETC (Senneff & Lowery, 2022):

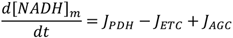

ETC flux (J_ETC_) is a simplified representation of the respiratory flux. J_ETC_ is driven by NADH_m_, with Michaelis constant q_3_ for NADH consumption, and is in inhibited at high ΔΨ through voltage dependence coefficients q_4_ and q_5_ (Senneff & Lowery, 2022):

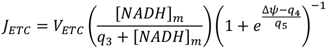

ATP synthesis is driven by the F1F0-ATPase flux (J_ATPase_), with maximal rate V_F1F0_. J_ATPase_ is inhibited at high mitochondrial ATP concentrations, with threshold value q_6_, and exhibits voltage dependence through coefficients q_7_ and q_8_ (Senneff & Lowery, 2022) :

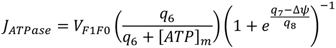

The adenine nucleotide translocator (J_ANT_) exchanges mitochondrial ATP for cytosolic ADP, with flux determined by the cytosolic and mitochondrial ADP/ATP ratios and ΔΨ (Senneff & Lowery, 2022):

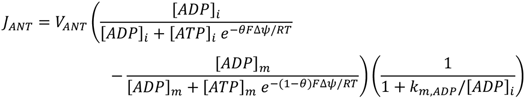

V_ANT_ is the maximal transport rate, *θ* is an empirical parameter, and k_m,ADP_ is a Michaelis constant.

Mitochondrial ATP and ADP concentrations are governed by the balance between ATP synthesis via J_ATPase_ and ATP export via J_ANT_ (Senneff & Lowery, 2022):

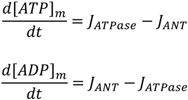

The Zukowski ECM-ROS model we assumed constant cytosolic [ATP_i_] and [ADP_i_] at 7.7mM and 0.29mM respectively (Cortassa *et al*., 2003), although these can be readily modified for future prediction. Recent experimental measurements using fluorescent ATP reporters in ventricular myocytes suggest that free cytosolic ATP concentrations may be substantially lower, below 1mM, than previously estimated from whole-heart preparations and fluctuate on a beat-to-beat timescale coupled to SR Ca^2+^ release and mitochondrial Ca^2+^ uptake (Rhana *et al*., 2024). The cytosolic ATP and ADP values derived from whole-heart preparations included here may be a future model extension.

Finally, proton leak across the inner mitochondrial membrane (J_HLeak_) is modeled as a linear function of ΔΨ (Senneff & Lowery, 2022):

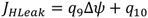

Where q_9_ is the voltage-dependent proton leak coefficient and q_10_ is the voltage-independent proton leak rate constant.

The mitochondrial membrane potential (ΔΨ) is governed by the balance of proton pumping via J_ETC_, ATP synthesis via J_ATPase_, adenine nucleotide exchange via J_ANT_, proton leak via J_HLeak_, and calcium flux contributions from J_NCX_, J_MCU_, J_mPTP_, and J_AGC_ (Senneff & Lowery, 2022):

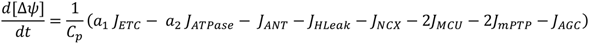

Where C_p_ is the mitochondrial inner membrane capacitance divided by Faraday’s constant (F), a_1_ is the scaling factor between NADH consumption and membrane voltage, and a_2_ is the scaling factor between ATP production by ATPase and change in membrane voltage.

### Model Coupling

To incorporate mitochondrial calcium fluxes into whole-cell calcium dynamics, the cytosolic calcium ODE of the ORd model was modified to include mitochondrial calcium uptake and efflux terms:

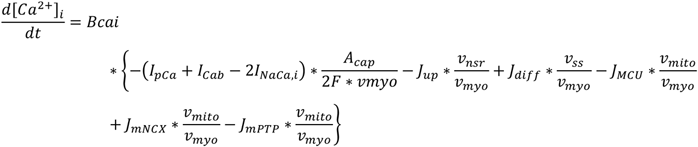

Where Bcai is the cytosolic calcium buffering factor, A_cap_ is the capacitive membrane area, F is Faraday’s constant, and the mitochondrial fluxes (J_MCU_, J_mNCX_, J_mPTP_) are scaled by v_mito_/v_myo_ to account for the difference in compartment volumes. The mitochondrial volume (v_mito_) is defined as 0.26*v_cell_. All other ORd calcium equations (subspace, JSR, NSR) were retained without modification.

### Reactive Oxygen Species (ROS) Module

The ROS module was adapted from the ROS-induced ROS release (RIRR) framework of Cortassa et al. (2004), which extends the integrated cardiac mitochondrial energetics model of Cortassa et al. (2003) to include mitochondrial ROS production, transport across the inner mitochondrial membrane, and cytosolic scavenging. The module is described by four ordinary differential equations governing mitochondrial superoxide ([O_2_ ^.-^]_m_), cytosolic superoxide ([O_2_ ^.-^]_i_), hydrogen peroxide ([H_2_O_2_]), and reduced glutathione ([GSH]) (Cortassa *et al*., 2004).

### Mitochondrial ROS Production

Mitochondrial superoxide ([O_2_^.-^]_m_) is produced as a fractional leak of electron transport chain flux which represents the proportion of J_ETC_ diverted to superoxide generation rather than proton pumping (Cortassa *et al*., 2004):

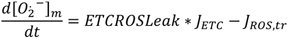

The ETC ROS leak fraction parameterizes the rate of mitochondrial superoxide production and serves as the primary determinant of steady-state ROS levels in the model. Mitochondrial superoxide is transported to the cytosol through the inner mitochondrial membrane anion channel (IMAC). The total IMAC conductance (J_IMAC_) is driven by ΔΨ and modulated by cytosolic superoxide concentration, which activates IMAC open probability, central to the RIRR framework (Cortassa *et al*., 2004). G_L_ is the leak conductance of IMAC, G_max_ is the integral conductance of IMAC at saturation, *κ* is a steepness factor for IMAC voltage dependence, and E_m,ROS_ is the membrane potential at half-saturation.

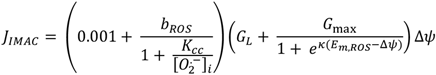

The superoxide-specific transport flux (J_ROS,tr_) represents the fraction of total IMAC current that carries superoxide across the inner membrane, scaled by the electrochemical driving force for superoxide transport (Cortassa *et al*., 2004):

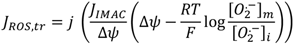

Where j is the fraction of IMAC conductance attributed to superoxide transport.

### Cytosolic ROS Dynamics

Cytosolic superoxide ([O_2_ ^.-^]_i_) accumulates through IMAC-mediated transport from the mitochondrial matrix and is scavenged by superoxide dismutase (SOD), which catalyzes its conversion to hydrogen peroxide ([H_2_O_2_]) (Cortassa *et al*., 2004):

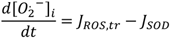

SOD activity (J_SOD_) incorporates the oxidized 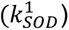, reduced 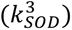, and inactive to active 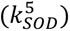 rate constants of the enzyme, with inhibition by H_2_O_2_ at high concentrations through inhibition constant 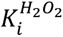:

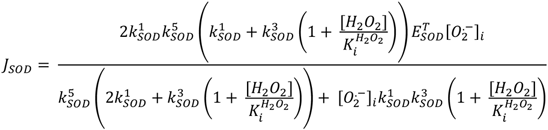

### Hydrogen Peroxide Scavenging and Glutathione Dynamics

Hydrogen peroxide ([H_2_O_2_]) accumulates through SOD activity and is scavenged by catalase (CAT) and glutathione peroxidase (GPx) (Cortassa *et al*., 2004):

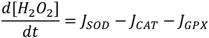

Catalase activity (J_CAT_) follows a first-order rate law with rate constant 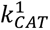, total catalase concentration 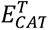, and an exponential inhibition factor capturing enzyme inactivation at high H_2_O_2_ concentrations (*fr*) (Cortassa *et al*., 2004):

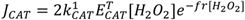

GPx activity (J_GPX_) depends on both H_2_O_2_ and reduced glutathione ([GSH]) concentrations, activity constants Φ_1_and Φ_2_, and intracellular GPX concentration 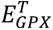 (Cortassa *et al*., 2004):

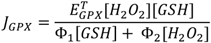

Reduced glutathione ([GSH]) is consumed by GPx and regenerated by glutathione reductase (GR), which uses NADPH as a cofactor. Cytosolic NADPH was held constant at 1mM consistent with Cortassa et al. (2004):

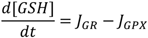

GR activity (J_GR_) is defined as:

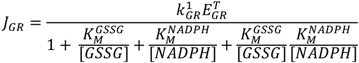

Where 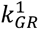 is the rate constant of GR, 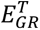 is the intracellular GR concentration, 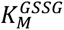 is the GSSG Michaelis constant and 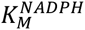 is the NADPH Michaelis constant. The total pool of glutathione, G_T_, is conserved:

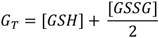

### ROS Dependent Ion Channel and Calcium Handling Modulation

Elevated cytosolic ROS modulates cardiac electrophysiology and calcium handling through both direct oxidative modification of ion channels and calcium handling proteins, and through Ca^2+^-independent oxidative activation of calcium/calmodulin-dependent protein kinase II (CaMKII) via modification of methionine residues M281/282 (Erickson *et al*., 2008). Oxidized CaMKII hyperphosphorylates the ryanodine receptor 2 (RyR2), increases sarcoplasmic reticulum Ca^2+^ leak, and augments the late sodium current (I_NaL_) (Wagner *et al*., 2011). Both superoxide and H_2_O_2_ increase RyR2 open probability in a concentration-dependent manner through oxidative modification of cysteine thiol groups (D’Oria *et al*., 2020). ROS additionally modulates sarcoplasmic reticulum Ca^2+^-ATPase (SERCA2a) activity through oxidative modification of phospholamban (PLN), which alters the PLN-SERCA2a inhibitory interaction (D’Oria *et al*., 2020).

ROS-dependent enhancement of the L-type calcium channel (I_CaL_) has also been demonstrated experimentally (Goldhaber & Liu, 1994; Hudasek *et al*., 2004; Hool & Corry, 2007), and augmentation of I_NaL_ under oxidative stress has been similarly characterized (Ward & Giles, 1997; Song *et al*., 2006). While the ORd model includes a simplified CaMKII phosphorylation term, it does not incorporate the detailed CaMKII activation dynamics necessary to mechanistically link cytosolic ROS to downstream channel phosphorylation. Therefore, in the Zukowski ECM-ROS model, ROS-dependent modulation was applied directly to RyR, SERCA, I_NaL_, and I_CaL_ as functions of cytosolic superoxide concentration ([O_2_ ^.-^]_i_) to simulate the experimentally reported net effect of oxidative stress on each target without explicit resolution of the intermediate CaMKII signaling cascade. A more detailed representation of ROS-CaMKII-channel coupling would require incorporation of explicit CaMKII activation kinetics that may be a future extension of the model.

Calibration of ROS-dependent scaling functions was performed by mapping experimental oxidative stress conditions to ROS concentration in the Zukowski ECM-ROS model. Experimental measurements used to calibrate ROS-dependent channel modulation were performed using exogenous H_2_O_2_ as a surrogate for intracellular oxidative stress. H_2_O_2_ is widely used in cardiac electrophysiology experiments because of its membrane permeability and relatively long half-life compared to superoxide. However, the doses used (often hundreds of micromolar) are beyond the range of physiological oxygen concentrations, and rapid intracellular catalase activity means that the effective intracellular concentration may not correspond to the applied extracellular dose (Ransy *et al*., 2020). In the Zukowski ECM-ROS model, scaling functions were applied as a function of cytosolic superoxide ([O_2_ ^.-^]_i_) consistent with the RIRR computational framework (Cortassa *et al*., 2004; Li *et al*., 2015), in which mitochondrial-derived superoxide is the primary ROS species driving downstream channel modulation. The application of H_2_O_2_-derived calibration data to superoxide-dependent scaling function is a first approximation modeling assumption. Future model refinements can include species-specific ROS scaling as experimental measurements of superoxide-dependent channel modulation become available.

### ROS-Dependent Modulation of Calcium Handling

Cytosolic ROS was coupled to sarcoplasmic reticulum calcium cycling through ROS-dependent scaling functions applied to ryanodine receptor release flux (J_RyR_) and SERCA uptake flux (J_up_). Scaling functions were derived from published experimental measurements of oxidative modulation of RyR open probability and SERCA activity, normalized to no ROS baseline conditions.

Redox modulation of ryanodine receptor (RyR)-mediated sarcoplasmic reticulum Ca^2+^ release was implemented using a data-driven scaling function derived from published single-channel RyR measurements. Experimental data relating H_2_O_2_ exposure to RyR open probability (P_o_) were obtained from Oba et al. (2002), in which skeletal muscle single-channel P_o_ was measured in lipid bilayers at defined redox potentials (cis = −220 mV, trans = −180 mV) across H_2_O_2_ concentrations of ≤10 µM (Oba *et al*., 2002). The Oba et al. (2002) dataset was selected because it reflects dose-dependent RyR activation data at low oxidant concentrations corresponding to the physiologically relevant ROS range of interest (10^-4^-10^-2^ mM; Sinenko *et al*., 2021). It should be noted that Oba et al. (2002) characterized skeletal muscle RyR1 rather than cardiac RyR2; cardiac RyR2-specific concentration-response data at comparable low ROS concentrations were not available at the time of model development, and the Oba et al. (2002) dataset was therefore used as the best available approximation of ROS-dependent RyR activation within the physiologically relevant concentration range.

Experimental data were digitized and normalized to baseline P_o_ to obtain a dimensionless fold-change in RyR activity as a function of oxidant concentration. Because the ORd model does not explicitly resolve RyR open probability, fold-change was applied multiplicatively to the RyR-mediated Ca^2+^ release flux (J_rel_). Normalized data were fit with a monotonically increasing exponential saturation function of the form:

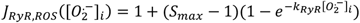

Where 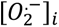 is cytosolic superoxide concentration (mM), *S_max_* represents the maximal fold-increase in RyR activity, and *k_RyR_* (mM^-1^) controls the steepness of the response. Nonlinear least-squares fitting yielded *S_max_* = 8.32 and *k_RyR_* = 27.01 *mM*^−1^ and captured the experimentally observed rapid increase and saturation of RyR activation with increasing oxidative stress. In simulations, cytosolic superoxide levels remain well below the saturation region of the fitted curve.

ROS inhibits SR Ca^2+^ uptake by impairing SERCA activity in a concentration-dependent manner (Xu *et al*., 1997). For SERCA uptake flux (J_up_), ROS-dependent inhibition of SERCA activity was characterized using data from Xu et al. (1997), in which purified cardiac SR vesicles prepared from rabbit hearts were exposed to a hydroxyl radical-generating system (H_2_O_2_ + Fe^3+^-NTA) at varying concentrations, and SERCA enzymatic activity was measured as a percent of control. Experimental data were digitized from Figure 1 of Xu et al. (1997) and normalized to baseline SERCA activity under no ROS conditions. Consistent with prior modeling work, redox-dependent SERCA inhibition was implemented as a multiplicative scaling of SERCA-mediated Ca^2+^ uptake flux (J_up_) using an exponential decay formulation (Li *et al*., 2015):

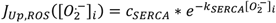

**Figure 1:**
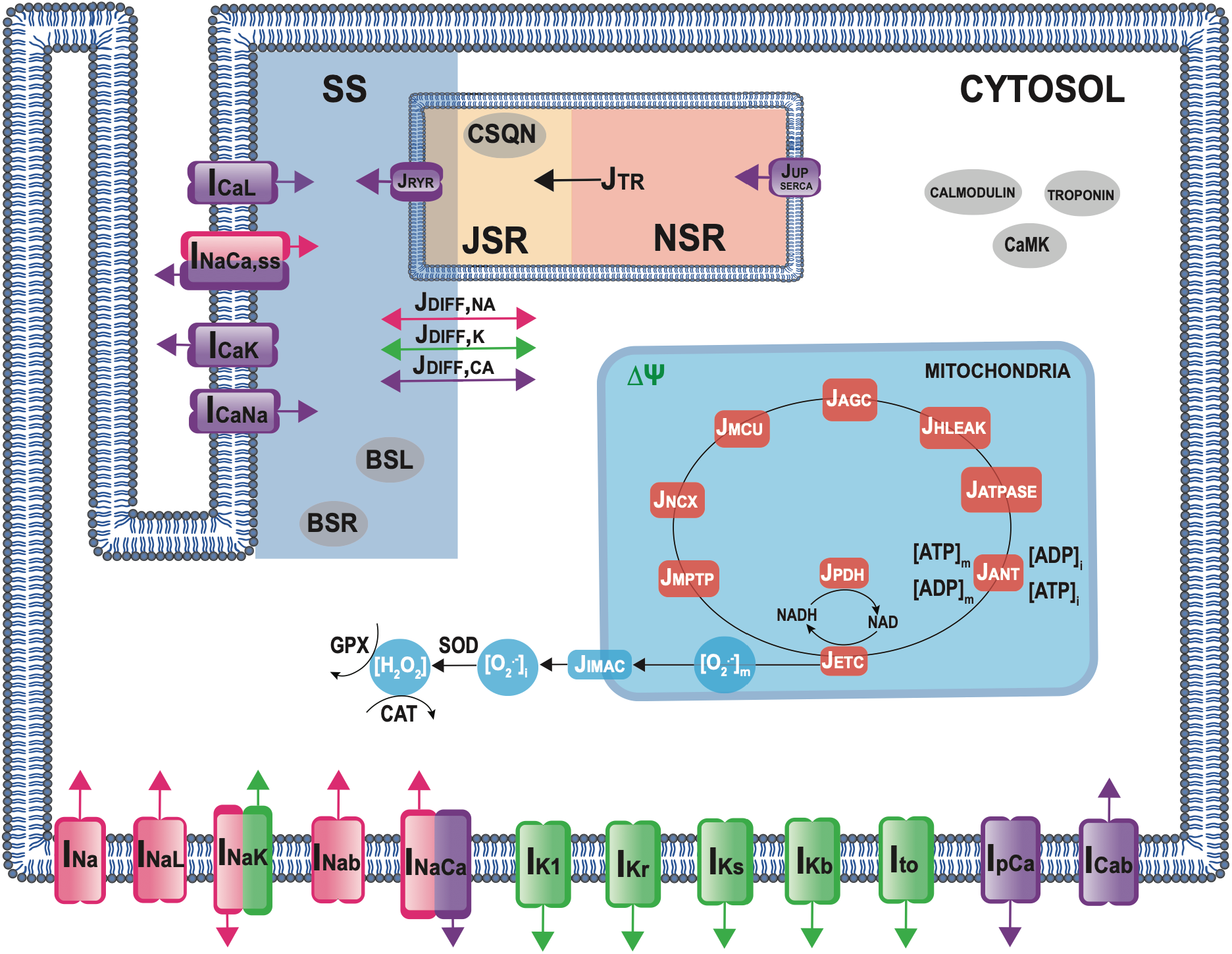
Schematic diagram of the Zukowski ECM-ROS computational model integrating human ventricular myocyte electrophysiology, mitochondrial energetics, and oxidative stress. The electrophysiology model derives from the ORd ventricular myocyte formulation (O’Hara *et al*., 2011), which includes membrane currents, intracellular ion diffusion, sarcoplasmic reticulum (SR) calcium fluxes, and buffering. Transmembrane currents (denoted by “I”) mediate the movement of ions as indicated by associated colors: calcium (purple), sodium (pink), and potassium (green). “SS” denotes the subspace compartment, “CYTOSOL” refers to the remaining intracellular space, “JSR” and “NSR” denote the junctional and network SR compartments, respectively. Fluxes are indicated by “J”. Mitochondrial fluxes appear in orange, and ROS generation and scavenging processes are shown in light blue. The Zukowski ECM-ROS model captures key processes underlying calcium-regulated energetics and is based on the Senneff and Lowery 2022 model (Senneff & Lowery, 2022). Mitochondrial calcium handling is mediated by the mitochondrial calcium uniporter (J_MCU_), sodium/calcium exchanger (J_NCX_), and mitochondrial permeability transition pore (J_mPTP_). Calcium activates the mitochondrial aspartate-glutamate carrier (J_AGC_), initiating NADH production via pyruvate dehydrogenase (J_PDH_) complex. The electron transport chain (J_ETC_) drives proton pumping and establishes the membrane potential (ΔΨ). ATP production is governed by F1F0-ATPase (J_ATPase_), and ATP export occurs through the adenine nucleotide translocator (J_ANT_), dependent on the cytosolic ATP/ADP ratio. A proton leak current (J_HLeak_) stabilizes ΔΨ. Reactive oxygen species (ROS) dynamics was built on the framework of Cortassa et al. (2004). Mitochondrial superoxide 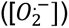 is produced as fractional leak of the electron transport chain (ETC) flux and is exported to the cytosol via the inner mitochondrial anion channel (J_IMAC_), driven by ΔΨ and the mitochondrial-to-cytosolic 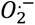 concentration gradient. In the cytosol, 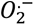 is rapidly converted into hydrogen peroxide ([H_2_O_2_]) by superoxide dismutase (SOD). H_2_O_2_ is scavenged through catalase (CAT) and glutathione peroxidase (GPx).

Where 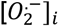 (mM) denotes cytosolic superoxide concentration, *c_SERCA_* is an inhibition coefficient, and *k_SERCA_* (mM^-1^) controls the steepness of the ROS-dependent inhibition. Nonlinear least-squares fitting yielded *c_SERCA_* = 1.02 and *k_SERCA_* = 43.67 *mM*^−1^ which reproduced experimentally observed suppression of SERCA function with increasing ROS.

Within the Zukowski ECM-ROS model, ROS-dependent scaling was applied multiplicatively to the baseline flux at each time step:

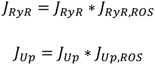

Under no ROS conditions ([O ^.-^]_i_ = 0), baseline calcium handling dynamics are recovered (J_RyR,ROS_ =1.0 and J_up,ROS_=1.02). Within increasing ROS, RyR Ca^2+^ release is progressively enhanced and SERCA Ca^2+^ uptake is progressively inhibited.

Both Xu et al. (1997) and Oba et al. (2002) applied exogenous oxidants under controlled *in vitro* conditions at concentrations that do not necessarily correspond to intracellular ROS concentrations under physiological or pathological conditions (Ransy *et al*., 2020). The scaling functions therefore represent an approximation of ROS-dependent RyR and SERCA modulation within the physiologically relevant ROS range (10^-4^-10^-2^ mM; Sinenko *et al.,* 2021), subject to the limitations of H_2_O_2_-based experimental techniques.

### ROS-Dependent Modulation of Ion Channels

ROS-dependent modulation of the late sodium current (I_NaL_) and L-type calcium current (I_CaL_) was implemented through saturating conductance scaling functions applied to the late sodium conductance (G_NaL_) and the L-type calcium channel phosphorylated permeability parameter (P_CaP_), respectively. P_CaP_ was selected as the ROS-dependent scaling target for I_CaL_, rather than baseline permeability (P_Ca_), because ROS are known to directly oxidize regulatory domain methionine residues of CaMKII (Erickson *et al*., 2008). The phosphorylated fraction of I_CaL_ in the ORd model is governed by CaMKII activity. Scaling P_CaP_ represents ROS-induced I_CaL_ enhancement acting through the oxidative CaMKII pathway. Experimental studies reporting oxidative modulation of I_NaL_ and I_CaL_ apply exogenous H_2_O_2_ in the micromolar to millimolar range, several orders of magnitude above intracellular [H_2_O_2_] under physiological or pathological conditions. Bath-applied [H_2_O_2_] do not directly reflect intracellular [ROS] (Ransy *et al*., 2020), and even modest elevation of intracellular H_2_O_2_ above ∼100nM are sufficient to activate stress-response signaling pathways and promote nonspecific oxidative modification (Chance, 1979; Sinenko *et al*., 2021). For these reasons, the reported experimental effects were interpreted as the maximal channel response under severe oxidative stress.

Scaling functions were implemented as logistic functions of cytosolic superoxide concentration, structured to reflect three concentration-dependent regimes (Chance, 1979; Sinenko *et al*., 2021): baseline channel behavior at low [ROS] ([O_2_ ^.-^]_i_ ≤ 100nM), a concentration-dependent transition across moderate oxidative stress (100nM < [O_2_ ^.-^]_i_ < 1µM), and a saturating maximal channel response under severe oxidative stress ([O_2_ ^.-^]_i_ ≥ 1µM). The general form of the scaling function is:

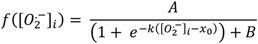

The offset term B was set such that f([O_2_ ^.-^]_i_ = 0) = 1.0 to ensure baseline channel behavior in the absence of ROS. The midpoint concentration x_0_ = 5.5 x 10^-4^ mM (550nM) was placed within the moderate-stress transition zone, and the steepness parameter k = 9000 mM^-1^ was chosen such that scaling remained near baseline through [O_2_ ^.-^]_i_ ≤ 100nM and approached its saturating maximum by [O_2_ ^.-^]_i_ = 1µM. The amplitude A was then determined by the constraint that f([O_2_ ^.-^]_i_ ≥ 1µM) equals the maximum scaling factor derived from experimental calibration, as follows.

For I_NaL_, Song et al. (2006) reported an increase of 81.78 ± 24.98% in I_NaL_ AUC (integral of I_NaL_) in isolated guinea pig ventricular myocytes exposed to 200µM H_2_O_2_ under a 300-ms voltage clamp protocol. Simulated I_NaL_ AUC at 1Hz was computed across a range of G_NaL_ scaling values until the percent increase fell within the experimentally reported range, which yielded a maximum G_NaL_ scaling factor of 2 at [O_2_ ^.-^]_i_ ≥ 1µM, with A = 1.0355 and B = 0.9823. For I_CaL_, Xie et al. (2009) reported an increase of 65.75 ± 30.63% in peak I_CaL_ in isolated adult rabbit ventricular myocytes exposed to 1mM H_2_O_2_ under a 300-ms voltage clamp protocol. Simulated peak I_CaL_ at 1Hz was computed at a range of P_CaP_ scaling values until the percent increase fell within the experimentally reported range, which yielded a maximum P_CaP_ scaling factor of 4 at [O_2_ ^.-^]_i_ ≥ 1 µM, with A = 3.1064 and B = 0.9468.

Scaling was applied within the ORd model ionic current formulation as:

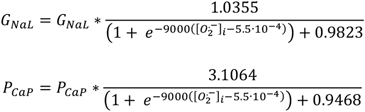

G_NaL_ and P_CaP_ scaling were calibrated simultaneously, as ROS-dependent enhancement of I_CaL_ and I_NaL_ interact through their effects on intracellular calcium and sodium dynamics. An increase in I_CaL_ scaling reduced the apparent I_NaL_ AUC increase, which required a tradeoff between the two calibration targets. The final parameter values represent the combination that placed both simulated I_NaL_ AUC and peak I_CaL_ increases within their respective experimental ranges simultaneously, with simulated peak I_CaL_ enhancement (45%) falling within the lower portion of the experimentally reported range (65.75 ± 30.63%; Xie *et al.,* 2009) and simulated I_NaL_ AUC enhancement (103%) falling within the experimentally reported range (81.78 ± 24.98%; Song *et al.,* 2006). Calibration results are presented in the Results section (**Figure 6**).

A key limitation of the available experimental data is that ROS concentration-dependent modulation of I_NaL_ and I_CaL_ across the full physiological ROS range are not available. The scaling functions therefore represent a first approximation based on a single maximum data point from each experimental study, with the concentration dependence governed by the functional form of the scaling function rather than direct experimental constraint. Future model refinements incorporating concentration-dependent experimental measurements of ROS-dependent I_NaL_ and I_CaL_ modulation would enable more physiologically grounded parameterization of scaling functions.

### Numerical Implementation

The model was implemented in C++ and integrated using the forward Euler method with a fixed time step of dt = 0.005ms. Population modeling and post-processing were performed in MATLAB 2025a. Cells were paced using a 0.5-ms, 80 µA/µF stimulus current at cycle lengths (CL) of 500, 750, 1000, and 2000ms and run to steady state (beat-to-beat changes in APD, peak [Ca^2+^]_i_, peak [O_2_^.-^]_i_, and peak [Ca^2+^]_m_ were < 1×10^-8^). Action potential duration was quantified at 50% (APD_50_) and 90% (APD_90_) repolarization. Calcium transients were quantified as peak amplitude of the steady-state transient. For calibration and validation simulations, cytosolic ROS concentration was held constant at specified values to reproduce experimental conditions. Model code, input parameter files, simulation scripts, and output files are publicly available on GitHub (https://github.com/ClancyLabUCD/A-Computational-Model-of-Oxidative-Stress-in-a-Human-Ventricular-Myocyte). The repository includes the C++ and MATLAB implementations of the Zukowski ECM-ROS model, MATLAB population modeling pipelines, and analysis scripts used to generate all figures reported in this study.

### Mitochondrial Calcium Validation

To validate model-predicted mitochondrial calcium uptake, simulations were performed under conditions matching Andrienko et al. (2009), in which steady-state mitochondrial calcium ([Ca^2+^]_m_) was measured as a function of fixed cytosolic calcium in single rat ventricular myocytes in the presence of 10mM Na^+^. Steady-state [Ca^2+^]_m_ dependence on [Ca^2+^]_i_ was extracted from Figure 3C of Andrienko et al. (2009). *In silico*, cytosolic calcium was fixed at values spanning the experimentally tested range (39.5, 156.9, 168.8, 300.1, 427.7, 553.4, 603.9, and 700µM), intracellular sodium was held constant at 10mM, and steady-state peak [Ca^2+^]_m_ was extracted from the final beat of a 5-beat simulation at 1Hz. Simulated values were compared against Andrienko et al. (2009) as the primary validation dataset, with data from table 1 of Collins et al. (2001) included as an independent comparison from a separate experimental preparation.

### APD_50_ Validation

To validate model-predicted ROS-dependent APD prolongation, cytosolic ROS ([O ^.-^] ) was held constant at 200µM and the model was paced for 50 beats at cycle lengths of 2000, 1000, 750, and 500ms (0.5–2Hz). Baseline simulations were performed with [O ^.-^] held constant at 1×10^-6^ mM. APD_50_ was extracted from the final beat and expressed as percent increase from baseline APD_50_. Simulated values were compared against experimental measurements from Song et al. (2006), in which guinea pig ventricular myocytes exposed to 200µM H_2_O_2_ with action potentials induced by a 5-ms depolarizing pulse at 0.16Hz exhibited 37.25 ± 13.41% APD_50_ prolongation relative to control, extracted from Figure 2 of Song et al. (2006). The experimental pacing rate (0.16Hz) falls outside the simulated range (0.5–2Hz), which was chosen to reflect physiologically relevant human heart rates from bradycardic to tachycardic conditions.

**Figure 2:**
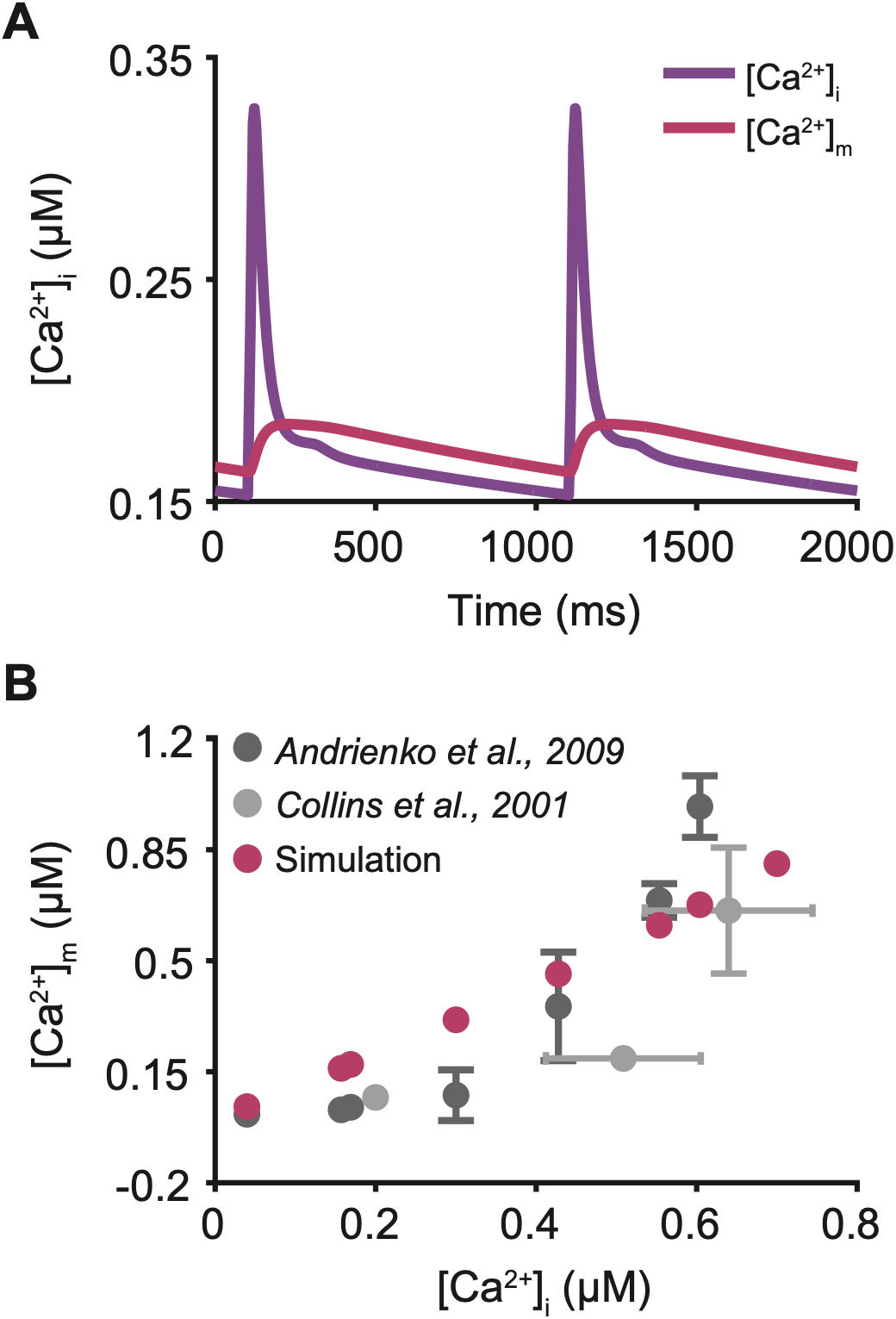
Prediction and validation of mitochondrial calcium dynamics in the Zukowski ECM-ROS model. **(A)** Simulated mitochondrial [Ca^2+^]_m_ (pink) and cytosolic [Ca^2+^]_i_ (purple) transients during steady-state 1.0Hz pacing. The synchronized oscillations between compartments are consistent with experimental observations reported by Chacon et al., 1996 (Chacon *et al*., 1996). **(B)** Steady-state peak mitochondrial [Ca^2+^]_m_ concentration (pink) as a function of fixed cytosolic [Ca^2+^]_i_. Experimental data from Andrienko et al., 2009 (dark grey) and Collins et al., 2001 (light grey) were used to validate model predictions under the same conditions. Error bars in experiments indicated standard deviation from the mean.

**Figure 3:**
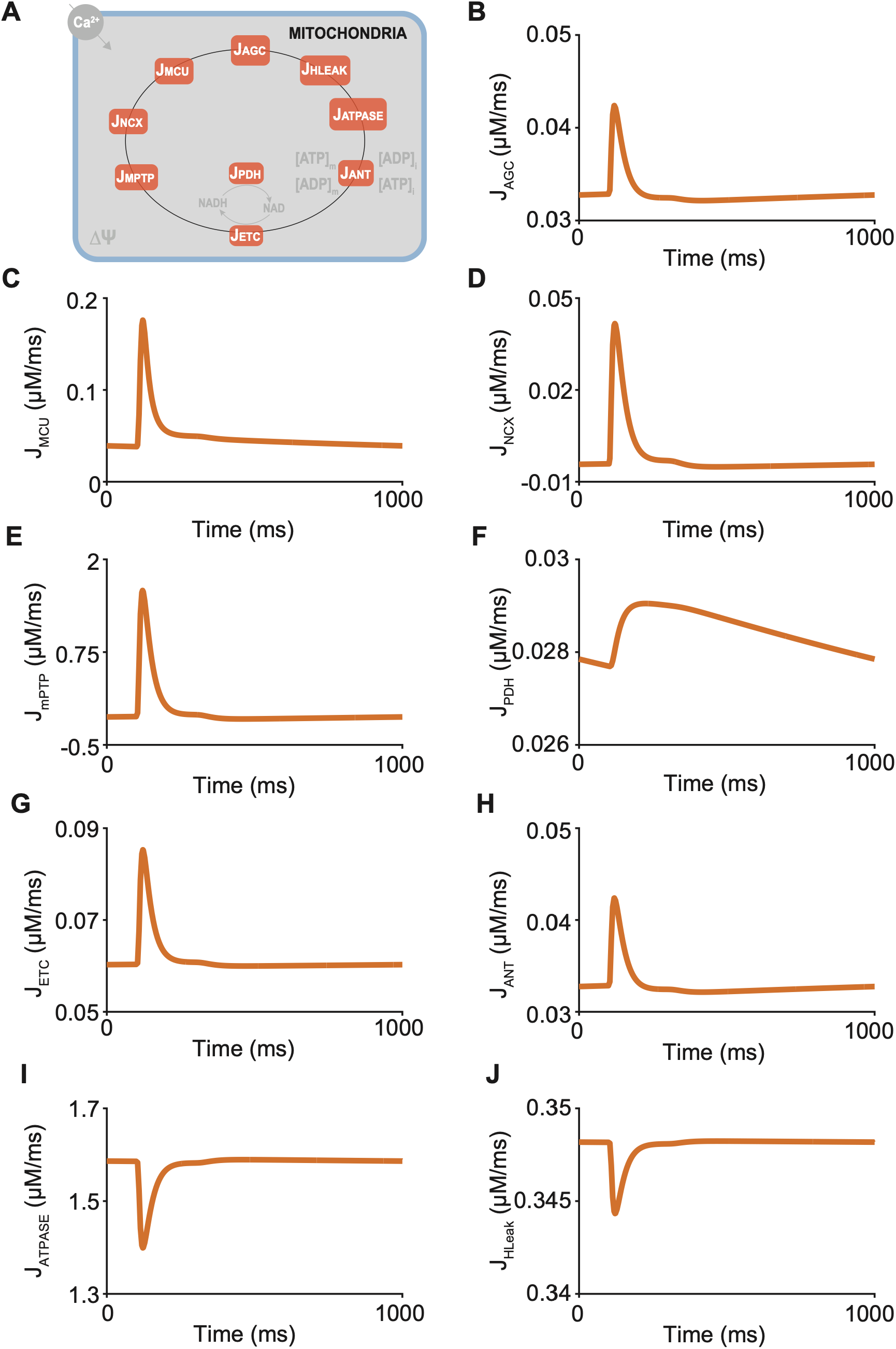
Steady-state mitochondrial flux dynamics in the Zukowski ECM-ROS model. Simulations were performed at steady state (CL=1000ms). **(A)** Schematic of the mitochondrial module adapted from Senneff and Lowery (2022), illustrating the nine mitochondrial fluxes included in the model (orange): mitochondrial calcium uniporter (J_MCU_), mitochondrial Na+/Ca+ exchanger (J_NCX_), mitochondrial permeability transition pore (J_mPTP_), aspartate-glutamate carrier (J_AGC_), pyruvate dehydrogenase complex (J_PDH_), ATP synthase (J_ATPase_), and proton leak (J_HLeak_). **(B-J)** Steady-state time course traces of each mitochondrial flux over one pacing cycle: **(B)** J_AGC_, **(C)** J_MCU_, **(D)** J_NCX_, **(E)** J_mPTP_, **(F)** J_PDH_, **(G)** J_ETC_, **(H)** J_ANT_, **(I)** J_ATPase_, and **(J)** J_HLeak_.

### Cytosolic Calcium Validation

To validate model-predicted ROS-dependent calcium handling, cytosolic ROS was held constant at 0.5mM and the model was paced for 50 beats at cycle lengths of 2000, 1000, 750, and 500ms. Baseline simulations were performed with [O_2_^.-^]_i_ held constant at 1×10^-6^ mM. Diastolic cytosolic calcium was calculated as percent increase from baseline at each pacing condition. The model’s diastolic (interbeat) [Ca^2+^]_i_ during pacing served as the closest available correlate to unpaced resting [Ca^2+^]_i_ to match experimental conditions. Simulated values were compared against experimental measurements from Wang et al. (1999), in which resting [Ca^2+^]_i_ in cell suspensions of Fura-2-loaded adult male Sprague-Dawley rat ventricular myocytes increased by 52.8 ± 4.7% following a 10-minute incubation with 0.5 mmol/L H_2_O_2_ at room temperature, extracted from Figure 3 of Wang et al. (1999).

### Cytosolic Calcium Perturbation Simulation Protocol

To investigate the model-predicted relationship between cytosolic calcium and mitochondrial energetics and ROS production, cytosolic and subspace calcium ([Ca^2+^]_i_, [Ca^2+^]_ss_) were scaled by a fixed multiplicative factor applied directly to the calcium concentration derivative:

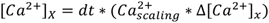

Where x = i (cytosol) or SS (subspace). 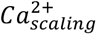 is the fixed scaling factor. Two sets of simulations were performed. First, at 1000ms cycle length, cytosolic and subspace calcium were scaled by factors of 1.3 and 1.5, which predicted steady-state peak cytosolic calcium increases of 37% and 62%, respectively. Time-dependent percent change from baseline was recorded for mitochondrial calcium ([Ca^2+^]_m_), electron transport chain flux (J_ETC_), mitochondrial membrane potential (ΔΨ), and cytosolic superoxide ([O_2_ ^.-^]_i_ ) following the calcium perturbation. Second, to evaluate the dependence of the cytosolic calcium to ROS relationship on pacing rate, cytosolic and subspace calcium were scaled by factors of 1.1 to 1.5 (in increments of 0.1), corresponding to peak cytosolic calcium increases of approximately 10–50%, at cycle lengths of 2000, 1000, 750, and 500ms. For each combination of calcium scaling factor and cycle length, the model was run to steady state (see Numerical Implementation). Percent change in cytosolic superoxide ([O_2_ ^.-^]_i_ ) from baseline was calculated, and the ROS amplification ratio was defined as the percent change in [O_2_ ^.-^]_i_ divided by the percent change in [Ca^2+^] for each condition.

### ETC ROS Leak Simulation Protocol

To investigate the model-predicted effects of mitochondrial ROS production on electrophysiology and calcium dynamics, the ETC ROS leak parameter, representing the fraction of electron transport chain flux (J_ETC_) contributing to mitochondrial ROS production ([O_2_^.-^]_m_), was varied from 0.40 to 0.60 (increment 0.01) at cycle lengths of 2000, 1000, 750, and 500ms. For each combination of ETC ROS leak and cycle length, the model was run to steady state (see Numerical Implementation), and APD_90_, peak cytosolic calcium ([Ca^2+^]_i_), peak mitochondrial calcium ([Ca^2+^]_m_), and peak cytosolic superoxide ([O ^.-^] ) were extracted. Representative time-domain traces of membrane voltage, cytosolic calcium, mitochondrial calcium, and cytosolic ROS were generated at a fixed ETC ROS leak of 0.55 across all cycle lengths to illustrate rate-dependent effects at a single representative leak value. The ETC ROS leak range (0.40–0.60) was selected to span a physiologically relevant range of cytosolic superoxide concentrations without producing unbounded accumulation, since values approaching the upper end of this range across all tested cycle lengths still allowed the model to reach a stable steady state under the convergence criteria described in **Numerical Implementation**.

### Doxorubicin (DOX) Parameterization

The Zukowski ECM-ROS computational model was applied to investigate doxorubicin (DOX)-induced cardiotoxicity. DOX is a widely used chemotherapy drug where cardiotoxic effects are driven by mitochondrial superoxide overproduction and impaired antioxidant scavenging (Benjanuwattra *et al*., 2020; Kong *et al*., 2022; Wu *et al*., 2022). DOX-induced cardiotoxicity is dose-dependent, with greater cumulative exposure associated with greater mitochondrial dysfunction and electrophysiological remodeling (Dudka *et al*., 2012; Li *et al*., 2025). To capture the dose dependence of DOX cardiotoxicity, DOX-induced stress was parameterized by a dimensionless scalar D (0 ≤ D ≤ 1), where D=0 represents baseline conditions and D=1 represents maximum DOX effect. D induces two independent perturbations to the mitochondrial ROS module: (1) enhanced ETC ROS leak and (2) reduced SOD scavenging capacity. ETC ROS leak and etSOD are the primary determinants of steady-state cytosolic superoxide ([O_2_ ^.-^]_i_ ), which is the species that directly modulates ion channel scaling functions. Other DOX-dependent effects, including direct ion channel modification and mitochondrial membrane damage, can be addressed in future model extensions.

In the Zukowski ECM-ROS computational model, “ETC ROS leak” parameter governs mitochondrial superoxide production, and the “etSOD” parameter governs the rate of cytosolic superoxide dismutation to H_2_O_2_. At baseline, (D=0, ETC ROS Leak = 0.35, etSOD = 1.5 x 10^-3^mM), the model predicts 1Hz steady-state peak [O_2_^.-^]_i_ = 61.8nM and [H_2_O_2_] =12.2nM, consistent with reported physiological ROS concentrations in healthy cardiomyocytes (Morawietz, 2023). Because ROS-dependent ion channel scaling functions (I_NaL_, I_CaL_) activate above 0.1µM [O_2_^.-^]_i_, there are no oxidative channel modifications at baseline (D=0).

Experimental studies demonstrate that DOX increases mitochondrial superoxide production at Complex I of the electron transport chain (Doroshow & Davies, 1986). DOX undergoes one-electron reduction at Complex I, forming a semiquinone radical that reduces molecular oxygen to superoxide while regenerating DOX. This process, known as redox cycling, diverts electron flux away from proton pumping and increases the fraction of superoxide production from the electron transport chain flux in proportion to DOX exposure (Marcillat *et al*., 1989). To model this mechanism, DOX-induced enhancement of mitochondrial superoxide production was implemented by scaling the ETC ROS leak parameter by a dimensionless multiplier α(D):

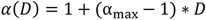

where α_max_ is the maximum ETC ROS leak scaling at full DOX stress (D=1). The scaled superoxide production term in the mitochondrial ROS equation becomes:

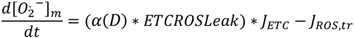

<u>_]_</u>Doroshow & Davies (1986) demonstrated that superoxide production increased approximately 40-fold at 90µM DOX in isolated beef heart submitochondrial particles. However, the 90µM DOX concentration used by Doroshow & Davies (1986) exceeds clinically relevant DOX concentrations. The precise magnitude of ETC ROS leak increase at clinically relevant intracellular DOX concentrations remains experimentally unconstrained. α_max_ was therefore set to 1.3 as a modeling parameter representing a ∼30% increase in the fraction of ETC electron flux leaking to superoxide at peak DOX stress (D=1), which corresponds to a maximum ETC ROS leak of 0.455. ETC ROS leak = 0.455 represents the upper boundary at which the model reaches steady state. ETC ROS leak values above 0.455, with DOX-induced decreased scavenging capacity, produce unbounded ROS accumulation due to the positive feedback between mitochondrial superoxide production and ROS-dependent channel modulation. Direct experimental quantification of ETC ROS leak changes at clinically relevant intracellular [DOX] will ideally be measured in future experiments.

In addition to enhanced superoxide production, DOX suppresses cellular antioxidant defense (Li & Singal, 2000). Doxorubicin treatment (cumulative dose 15mg/kg over 2 weeks) reduced GPx activity by up to 32% and transiently reduced mitochondrial SOD activity 20-27% within 2-24 hours of administration (Li & Singal, 2000). DOX-induced reduction in antioxidant scavenging capacity was implemented in the Zukowski ECM-ROS model by scaling the parameter “etSOD” by a dimensionless multiplier β(D):

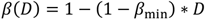

where *β*_min_ defines the minimum scavenging capacity at full DOX stress (D=1) and β(D=0) = 1.0 recovers baseline SOD scavenging capacity in the absence of DOX. The effective SOD concentration becomes:

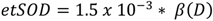

*β*_min_ was set to 0.75 which corresponds to a 25% reduction in scavenging capacity at D=1, consistent with the magnitude of acute antioxidant enzyme reductions reported experimentally (Li & Singal, 2000). Although the full ROS scavenging system in the model includes GPx, catalase, and glutathione reductase, DOX-induced antioxidant depletion was implemented exclusively through scaling of etSOD. Because electrophysiological remodeling in the model is driven by cytosolic superoxide rather than downstream H_2_O_2_ or glutathione species, etSOD (which governs the rate of cytosolic superoxide dismutation) is the primary scavenging parameter that modulates the ROS-calcium positive feedback loop in the model. Scaling etSOD by β(D) is therefore assumed to capture the net effect of DOX-induced antioxidant depletion on electrophysiologically relevant ROS (cytosolic superoxide, [O ^.-^] ). The contribution of DOX-dependent reductions in GPx, catalase, and glutathione reductase to cytosolic superoxide accumulation is a future modeling extension.

### DOX-Induced APD Prolongation Validation

To validate model-predicted DOX-induced APD prolongation, simulations were performed at 1Hz pacing across the full range of DOX severity levels (D=0 to D=1.0). Baseline steady-state APD_80_ was extracted at D=0, and percent change in APD_80_ was calculated at D=1.0 relative to baseline. Because the model’s DOX severity parameter (D) represents a generalized level of DOX-induced stress rather than a specific pharmacological concentration, simulated percent change in APD_80_ at maximal DOX effect (D=1.0) was compared against the full range of experimentally reported DOX concentrations, rather than against a single matched dose. Simulated values were compared against experimental data from George et al. (2026), in which left ventricular slices from young male human donors (n=9, age = 39 ± 9 years) were cultured in DOX at concentrations of 0.5, 1, 5, and 10µM for 24 hours, then optically mapped during 1Hz pacing to measure APD_80_, with APD_80_ change of 2-13% (SD range 7-20%) reported relative to control across this concentration range (George *et al*., 2026).

### DOX Dose-Response Simulation Protocol

To characterize DOX-induced mitochondrial and electrophysiological dysfunction across pacing rates, steady-state simulations were performed across DOX effect levels (D=0, 0.5, 0.6, 0.7, 0.75, 0.8, 0.9, 1.0) at cycle lengths of 500, 750, and 1000ms. For each combination of D and cycle length, peak cytosolic calcium ([Ca^2+^]_i_), peak mitochondrial calcium ([Ca^2+^]_m_), peak cytosolic superoxide ([O_2_ ^.-^]_i_ ), minimum mitochondrial membrane potential (ΔΨ), and APD were extracted. Percent change for each variable was calculated relative to baseline (D=0) at the corresponding cycle length.

To evaluate the temporal relationship between mitochondrial dysfunction and electrophysiological remodeling, the 500ms (2Hz) cycle length condition was examined separately across DOX effect levels D=0.5–1.0. Percent change from baseline (D=0) was compared across APD_90_, cytosolic superoxide ([O ^.-^] ), mitochondrial calcium ([Ca^2+^]_m_), and mitochondrial membrane potential (ΔΨ) to assess the relative onset and magnitude of mitochondrial versus electrophysiological changes.

#### Population Modeling

Cancer patients treated with DOX are frequently co-administered medications known to prolong the QT interval through block of the rapid delayed rectifier potassium current (I_Kr_) (Agnihotri *et al*., 2024; Ramasubbu *et al*., 2025). To investigate whether DOX-induced oxidative stress and pharmacological I_Kr_ block combine to increase arrhythmia susceptibility, five simulation conditions were examined: (1) no DOX (D=0), 0% I_Kr_ block; (2) full DOX effect (D=1), 0% I_Kr_ block; (3) full DOX effect (D=1), 30% I_Kr_ block; (4) full DOX effect (D=1), 40% I_Kr_ block; and (5) no DOX (D=0), 40% I_Kr_ block. Conditions 1 and 2 isolate the effect of DOX alone, conditions 3 and 4 examine DOX combined with progressive I_Kr_ block, and condition 5 isolates the effect of I_Kr_ block alone. I_Kr_ block was implemented as a fixed scaling constant on maximal I_Kr_ conductance (G_Kr_), hard-coded for each condition (IKr_scaling = 1.0, 0.7, or 0.6 for 0%, 30%, and 40% block, respectively).

A population of N=1000 virtual human ventricular cells were generated to investigate inter-individual variability in ion channel expression under DOX. Twelve ion channel conductance scaling factors (GNa, Gto, GNaL, PCa, GKs, GK1, GNCX, PNaK, GKb, PNab, PCab, GpCa; **Table 2**) were independently sampled from a uniform distribution spanning 0.8-1.2x (±20%) baseline for each cell to simulate physiological variability in channel expression across a virtual patient population. A fixed random seed was used to generate a single population of 1000 virtual cells, with the same twelve conductance scaling factors applied identically across all five simulation conditions to ensure the differences in outcomes between conditions reflect the applied DOX effect and I_Kr_ block perturbations rather than variability in the sampled population.

**Table 2:** Ion channel conductance parameters varied ±20% in population simulations. Each parameter was independently sampled from a uniform distribution within ±20% of its baseline value across N=1000 virtual cells to simulate inter-individual variability in ion channel expression.

| Ion Channel | Conductance Parameter Scaled $\pm 20\%$ |
| --- | --- |
| $I_{Na}$ | GNa |
| $I_{NaL}$ | GNaL |
| $I_{NaCa}$ | Gncx |
| $I_{NaK}$ | Pnak |
| $I_{Ks}$ | GKs |
| $I_{K1}$ | GK1 |
| $I_{to}$ | Gto |
| $I_{CaL}$ | Pca |
| $I_{Kb}$ | GKb |
| $I_{Nab}$ | PNab |
| $I_{Cab}$ | PCab |
| $I_{pCa}$ | GpCa |

For each of the five conditions, every cell in the population was simulated as a single continuous run: starting from fixed initial conditions representing 2Hz steady-state under that condition’s DOX effect (D=0 or D=1), each cell was paced at 2Hz (cycle length 500ms) for 10 beats, followed by a single 1000ms pause, after which one additional beat was recorded using forward Euler integration (dt=0.01ms). DOX effect and I_Kr_ block were applied throughout the full simulation for each condition. Pause-dependent early afterdepolarizations have been implicated in the initiation of torsade de pointes in the setting of prolonged repolarization and computational modeling has demonstrated that a pause following rapid pacing can unmask EAD susceptibility not evident during steady-state pacing alone (Viswanathan & Rudy, 1999; Liu & Laurita, 2005; Tang *et al*., 2012). A trial was flagged as failed and excluded from subsequent analysis if, at the end of the pacing protocol, cytosolic calcium was negative, cytosolic calcium exceeded 5.0mM, or membrane potential was non-finite. No trials met these exclusion criteria across the five simulation conditions.

For each cell, membrane potential, cytosolic calcium ([Ca^2+^]_i_), mitochondrial calcium ([Ca^2+^]_m_), and cytosolic ROS ([O_2_ ^.-^]_i_ ) traces were recorded for the post-pause beat. APD_90_ was calculated from the voltage traces. Mean and standard deviation of APD_90_ were calculated across the population of 1000 cells for each of the five simulation conditions.

## RESULTS

The Zukowski ECM-ROS model comprises a bidirectionally coupled model of excitation– contraction coupling, mitochondrial energetics, and redox signaling in the human ventricular myocyte. Zukowski ECM-ROS was developed to investigate interactions between cardiac electrophysiology, mitochondrial energetics, calcium handling, and reactive oxygen species (ROS) dynamics. The model integrates the O’Hara-Rudy (ORd) human ventricular myocyte model (O’Hara *et al*., 2011), mitochondrial energetics modified from the Senneff-Lowery model (Senneff & Lowery, 2022), and ROS production and scavenging from the Cortassa RIRR model (Cortassa *et al*., 2004) (**Fig. 1**).

Within the integrated framework, the membrane voltage and ionic currents are generated by the ORd model and regulate intracellular calcium cycling, which serves as the primary input to the mitochondrial system. Cytosolic calcium influences mitochondrial calcium uptake and metabolic activity through calcium-sensitive mitochondrial transport and energetic pathways. Mitochondrial activity regulates ROS production through electron transport chain ROS leak. Cytosolic ROS then feeds back onto whole-cell electrophysiology and calcium handling through redox-dependent modulation of ion channels (I_NaL_, I_CaL_) and sarcoplasmic reticulum calcium cycling (J_RyR_, J_up_).

The integrated model provides a framework for investigating how alterations in calcium handling and mitochondrial ROS production interact to influence cardiac electrophysiology across physiological and pathological conditions.

To validate the mitochondrial calcium handling predicted by the Zukowski ECM-ROS model, simulated mitochondrial calcium dynamics were compared to experimentally reported relationships between cytosolic and mitochondrial calcium. Under steady state 1Hz pacing conditions, the model predicted synchronized oscillations between cytosolic calcium ([Ca^2+^]_i_) and mitochondrial calcium ([Ca^2+^]_m_) (**Fig. 2A**). Simulated mitochondrial calcium transients followed the time course of the cytosolic calcium transient while maintaining smaller amplitude and slower dynamics, consistent with experimentally observed coupling between cytosolic and mitochondrial calcium compartments (Chacon *et al*., 1996).

To validate model-predicted mitochondrial calcium uptake against experimental measurements, simulations were performed under conditions reported by Andrienko et al. (2009). Cytosolic calcium concentration was fixed at values corresponding to the experimentally tested range and predicted steady-state peak mitochondrial calcium was plotted at each concentration (**Fig. 2B**). Simulated peak mitochondrial calcium was predicted to exhibit a positive relationship with cytosolic calcium concentration within the experimentally reported ranges (Collins *et al*., 2001; Andrienko *et al*., 2009). The close agreement between model-predicted and experimentally reported mitochondrial calcium concentrations demonstrates that the model accurately captures mitochondrial calcium uptake and buffering dynamics.

Within the Zukowski ECM-ROS model, cytosolic calcium ([Ca^2+^]_i_) serves as the primary input linking excitation-contraction coupling to mitochondrial calcium handling and metabolism. Cytosolic calcium enters the mitochondria through the mitochondrial calcium uniporter (J_MCU_) and is extruded through the mitochondrial Na^+^/Ca^2+^ exchanger (J_NCX_) and mitochondrial permeability transition pore (J_mPTP_). Mitochondrial calcium additionally regulates metabolic flux through activation of the aspartate-glutamate carrier (J_AGC_) and pyruvate dehydrogenase complex (J_PDH_), coupling beat-to-beat dynamics to NADH production and electron transport chain (J_ETC_) activity **(Fig. 3A)**. To characterize the steady-state behavior of the Zukowski ECM-ROS model and allow verification and reproducibility for future users, simulations were performed at pacing cycle length of 1000ms. Representative steady-state traces of all mitochondrial fluxes are shown in **Fig. 3B-J**. Each predicted flux exhibits stable beat-to-beat oscillations synchronized with the cytosolic calcium transient, consistent with physiological coupling between whole-cell electrophysiology and mitochondrial energetics.

To investigate the response of mitochondrial energetics and ROS production to altered cytosolic calcium dynamics, perturbations were applied in the Zukowski ECM-ROS model to the cytosolic and subspace calcium update terms (Δ[Ca^2+^]_i_ , Δ[Ca^2+^]_ss_) (**Fig. 4A**). Cytosolic calcium model dynamics were scaled by factors of 1.1–1.5 relative to baseline conditions (scaling factor = 1.0). Simulations in **Fig. 4A-E** were conducted at steady state 1Hz pacing with calcium scaling factors of 1.3 and 1.5, corresponding to predicted increases of 37% and 62% in peak [Ca^2+^]_i_, respectively. The predicted increase in cytosolic calcium was quantified as percent change relative to baseline steady-state calcium dynamics, and downstream changes in mitochondrial calcium ([Ca^2+^]_m_), mitochondrial membrane potential (ΔΨ), electron transport chain flux (J_ETC_), and cytosolic ROS ([O_2_ ^.-^]_i_) were subsequently calculated.

**Figure 4:**
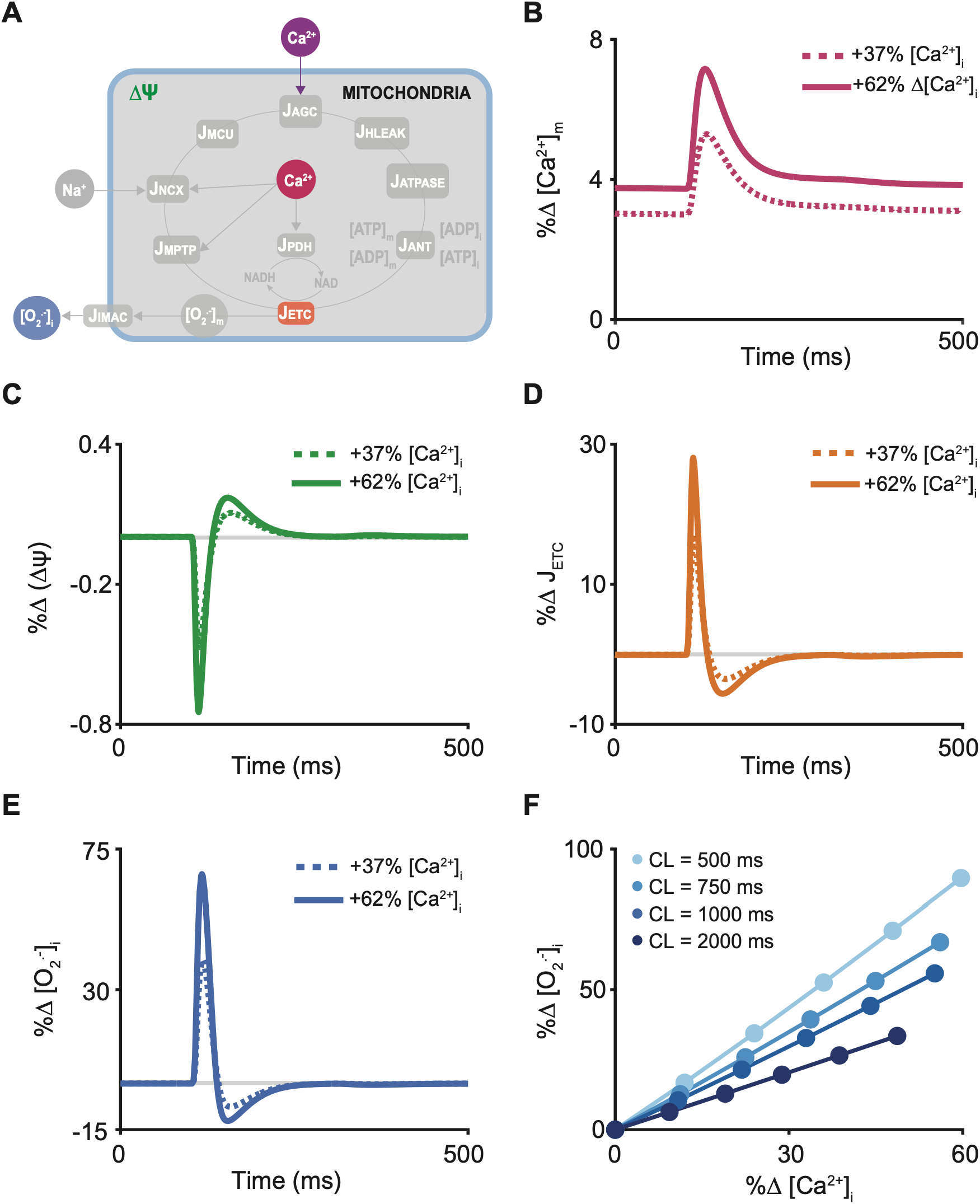
The predicted response of mitochondrial dynamics to increased peak cytosolic [Ca^2+^]_I_ in the Zukowski ECM-ROS model. Simulations in (A)-(E) were performed at steady state (CL = 1000ms). **(A)** A Schematic of the mitochondrial-ROS module is shown. Cytosolic calcium (Ca^2+^, purple) serves as the primary input to the mitochondria and links whole-cell electrophysiology to mitochondrial energetics and ROS production. Ca^2+^ influx through the mitochondrial calcium uniporter (J_MCU_) and efflux via mitochondrial sodium-calcium exchanger (J_NCX_) and mitochondrial permeability transition pore (J_mPTP_) regulate mitochondrial Ca^2+^ concentration (Ca^2+^, pink), which in turn modulates mitochondrial membrane potential (ΔΨ, green), electron transport chain flux (J_ETC_, orange), and cytosolic superoxide ([O ^.-^] , blue). Time-dependent percent change (Δ%) in key system variables follows a transient increase in peak cytosolic Ca^2+^ of 62% (solid lines) and 37% (dashed lines), relative to baseline. Cytosolic Ca^2+^ was scaled in cytosolic and subspace calcium (Δ[Ca^2+^]_i_ , Δ[Ca^2+^]_ss_ ) by factors of 1.3 and 1.5. Shown are predicted model responses to increased cytosolic and subspace calcium (Δ[Ca^2+^]_i_ , Δ[Ca^2+^]_ss_ ) in **(B)** mitochondrial Ca^2+^ transient ([Ca^2+^]_m_), **(C)** mitochondrial membrane potential (ΔΨ), **(D)** electron transport chain flux (J_ETC_), and **(E)** cytosolic superoxide production ([O ^.-^] ). **(F)** Relationship between percent change in cytosolic Ca^2+^ ([Ca^2+^]_i_) and percent change in cytosolic superoxide ([O ^.-^] ) across pacing cycle lengths (CL = 2000, 1000, 750, and 500ms). Cytosolic calcium dynamics were scaled by factors of 1.1-1.5 relative to baseline for cytosolic and subspace calcium (Δ[Ca^2+^]_i_, Δ[Ca^2+^]_ss_ ). The resulting changes in cytosolic calcium were quantified relative to baseline calcium and corresponding percent changes in [O .] were predicted.

Increased cytosolic calcium predicted progressive elevation of mitochondrial calcium, with peak [Ca^2+^]_m_ increasing 3.4% and 4.3% and diastolic [Ca^2+^]_m_ increasing 3.0% and 3.7% at 37% and 62% calcium perturbations respectively (**Fig. 4B**), consistent with enhanced calcium influx through the mitochondrial calcium uniporter. Elevated mitochondrial calcium was accompanied by increased electron transport chain flux, with peak J_ETC_ increasing 11.8% and 18.8% at 37% and 62% perturbations respectively (**Fig. 4C**), and mitochondrial membrane potential depolarization of 0.34% and 0.53% (**Fig. 4D**). Cytosolic ROS was predicted to have the largest percent changes of all examined variables, with peak [O_2_ ^.-^]_i_ increasing 32.0% and 52.6% at 37% and 62% calcium perturbations respectively (**Fig. 4E**), consistent with elevated ROS generation secondary to increased ETC activity. The disproportionate amplification of ROS relative to the modest changes in mitochondrial calcium and ETC flux reflects the nonlinear sensitivity of ROS production to mitochondrial calcium loading within the model.

To predict the relationship between cytosolic calcium and ROS production for a range of pacing frequencies, calcium increase of 10–50% (scaling factors 1.1-1.5) were applied at cycle lengths of 2000, 1000, 750, and 500ms (**Fig. 4F**). Across all pacing frequencies, increased cytosolic calcium was predicted to increase cytosolic ROS, with the magnitude of ROS amplification strongly dependent on pacing rate. At a 50% calcium increase, peak [O_2_^.-^]_i_ was predicted to increase 33.6% at CL=2000ms compared to 89.5% at CL=500ms which corresponds to a 2.66-fold greater ROS response at faster pacing. The ROS amplification ratio (%Δ[O_2_^.-^]_i_ / %Δ[Ca^2+^]_i_) increased from 0.69 at CL=2000ms to 1.50 at CL=500ms, indicating that at slower pacing rates a calcium increase produces a less than proportional ROS increase, while at faster pacing rates the same perturbation produces a greater than proportional ROS response. The consistent 2.66-fold difference in ROS amplification between CL=500ms and CL=2000ms for all increased calcium levels demonstrates that pacing rate exerts a multiplicative effect on ROS production independent of the increase in calcium. This prediction is consistent with the reduced diastolic recovery time at faster pacing rates, which promotes progressive mitochondrial calcium accumulation and increased electron transport chain (ETC) driven ROS generation.

To incorporate experimentally observed ROS-dependent modulation of sarcoplasmic reticulum (SR) calcium handling, cytosolic ROS was coupled to both ryanodine receptor (RyR) release flux (J_RyR_) and SERCA uptake flux (J_up_) in the Zukowski ECM-ROS model (**Fig. 5A**). ROS-dependent scaling functions were calculated from experimental measurements describing oxidative modulation of RyR open probability and SERCA activity (**Fig. 5B**). For RyR, fold-change in channel open probability was extracted from Oba et al. (2002), normalized to baseline RyR open probability in the absence of ROS, and fit with a sigmoidal function to define ROS-dependent modulation of J_RyR_. For SERCA, ROS-dependent changes in SERCA flux were extracted from Xu et al. (1997), normalized to baseline SERCA activity under no ROS conditions, and similarly fit with a sigmoidal function to define ROS-dependent modulation of J_up_. Together, these functions enabled ROS-dependent modulation of intracellular calcium cycling between the cytosol, network SR (NSR), junctional SR (JSR), and mitochondria.

**Figure 5.**
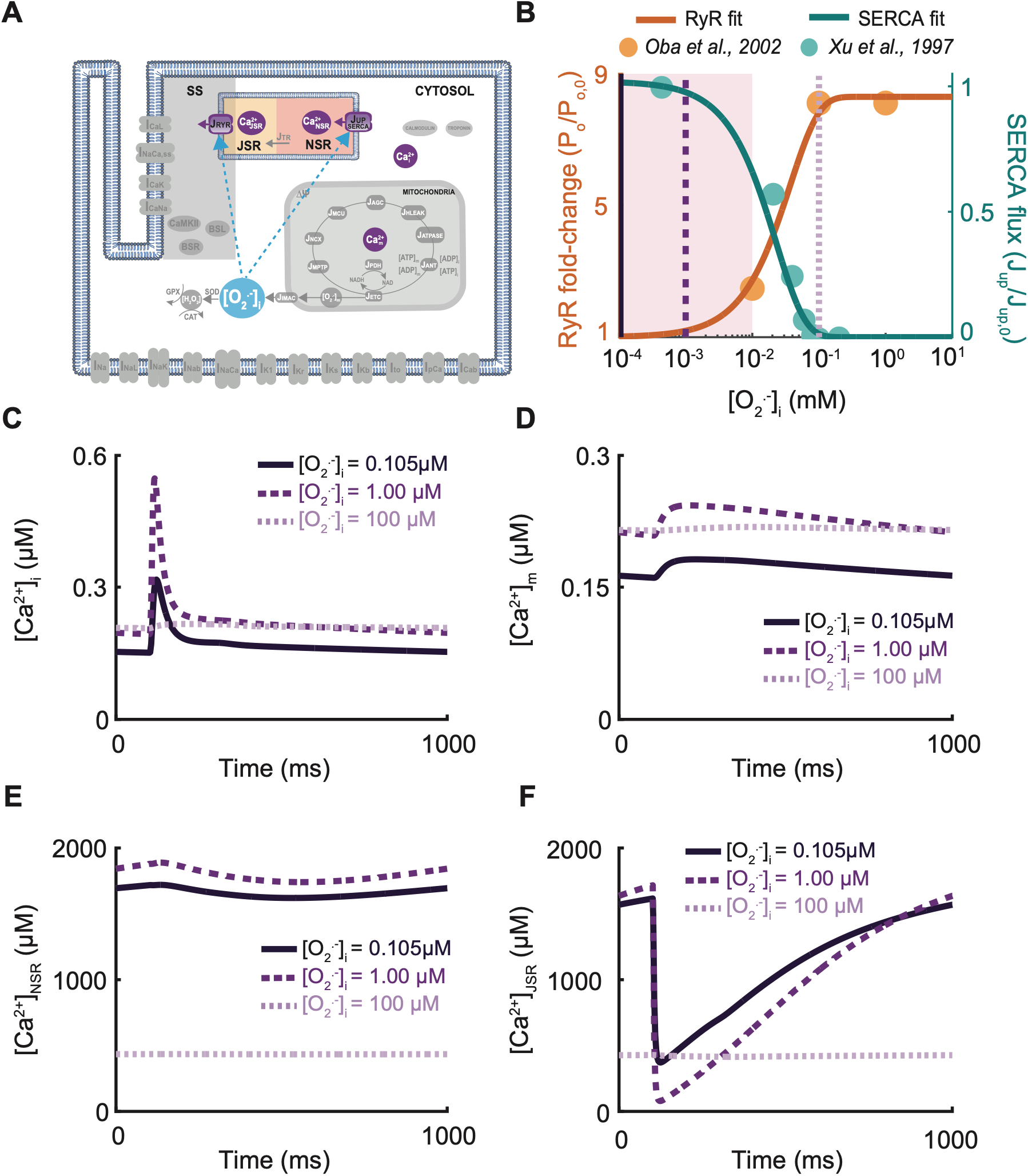
ROS-dependent modulation of sarcoplasmic reticulum Ca^2+^ handling is predicted to produce nonlinear effects on intracellular Ca^2+^ dynamics. **(A)** Schematic of the Zukowski ECM-ROS model highlighting cytosolic ROS effects on ryanodine receptor flux (J_RyR_), SERCA flux (J_Up_), and downstream calcium in the NSR, JSR, cytosol, and mitochondria. **(B)** A ROS-dependent scaling function based on experimental data (Xu *et al*., 1997; Oba *et al*., 2002) was applied to RyR (J_RyR_ = J_RyR_ * J_RyR,ROS_) and SERCA (J_SERCA_ = J_SERCA_ * J_SERCA,ROS_) to model redox modulation of Ca^2+^ handling. The shaded region represents relevant physiological ROS concentrations (10^-4^–10^-2^ mM) (Sinenko *et al*., 2021). Vertical dashed lines indicate representative ROS levels used for time-domain simulations in **(C)** cytosolic calcium, **(D)** mitochondrial calcium, **(E)** network sarcoplasmic reticulum (NSR) calcium, and **(F)** junctional sarcoplasmic reticulum (JSR) calcium. Simulations are shown at [O ^.-^] = 0.105µM (solid), 1.00µM (dashed), and 100µM (dotted). Moderate ROS enhanced cytosolic calcium transient amplitude via increased RyR-mediated release, while high ROS drove progressive NSR and JSR depletion, reflecting pathological SR calcium dysregulation under severe oxidative stress.

**Figure 6:**
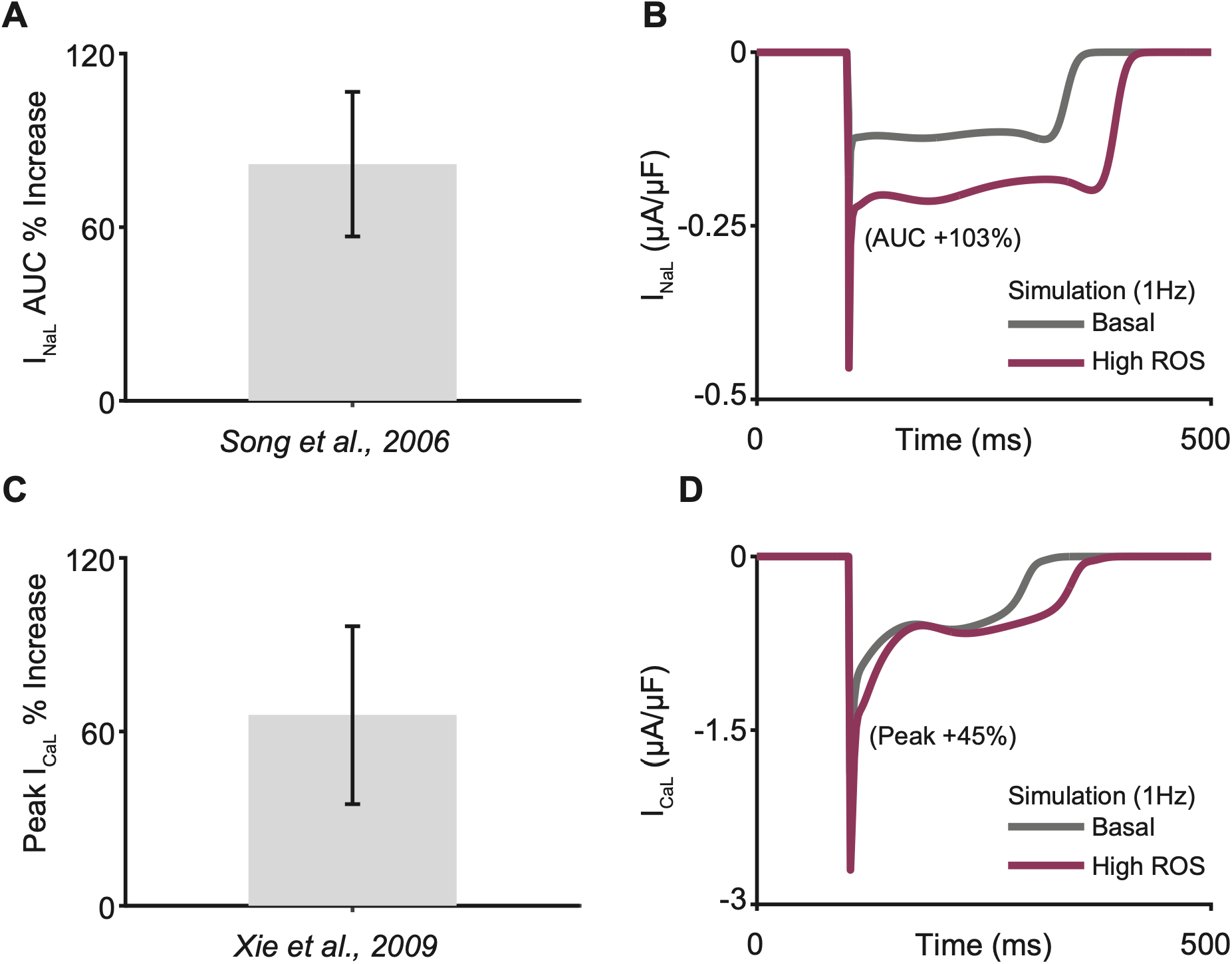
Zukowski ECM-ROS model calibration to experimental measurements of ROS-dependent modulation of late sodium and L-type calcium currents. **(A)** Percent increase in late sodium current (I_NaL_) area under the curve (AUC) under maximal ROS exposure. Experimental range (grey bar; Song *et al.,* 2006) reflects measurements in guinea pig ventricular myocytes exposed to 200µM H_2_O_2_ under a 300-ms voltage clamp protocol. I_NaL_ conductance was tuned in the model such that maximal I_NaL_ ROS-dependent enhancement at 1Hz falls within experimentally observed range. **(B)** Steady state 1Hz simulated I_NaL_ traces under basal (grey) and high ROS (maroon) conditions. High ROS increases I_NaL_ AUC by 103% compared to basal. **(C)** Percent increase in peak L-type calcium current (I_CaL_) under maximal ROS exposure. Experimental range (grey bar; Xie *et al.,* 2009) reflects measurements in adult rabbit ventricular myocytes exposed to 1mM H_2_O_2_ under a 300-ms voltage clamp protocol. I_CaL_ ROS-dependent conductance was tuned such that maximal I_CaL_ enhancement at 1Hz matched experimental conditions. **(D)** 1Hz steady state simulated I_CaL_ traces under basal (grey) and high ROS (maroon) conditions. High ROS increases peak I_CaL_ by 45% compared to basal. In all simulations, maximal ROS concentrations correspond to peak [O ^.-^] ≥100µM (Sinenko *et al*., 2021). Error bars in experiments indicate standard deviation from the mean of the experimental data.

The shaded region in **Fig. 5B** represents the physiologically relevant ROS concentration range investigated in this study (10^-4^–10^-2^ mM). Within this range, moderate ROS exposure (0.105µM and 1.00µM) was predicted to increase cytosolic calcium transient amplitude because of enhanced RyR-mediated calcium release (**Fig. 5C-D**). However, at higher ROS concentrations (100µM), the model predicted depletion of calcium from both the NSR and JSR, consistent with impaired SR calcium storage under severe oxidative stress (**Fig. 5E-F**). The transition from enhanced calcium release to SR calcium depletion is experimentally supported by observations of reduced SR calcium content following radiation-induced oxidative stress in ventricular myocytes (Sag *et al*., 2013).

Together, these results demonstrate that ROS-dependent modulation of RyR and SERCA produces concentration-dependent effects on intracellular calcium handling, with moderate ROS enhancing cytosolic calcium transients and high ROS driving pathological SR calcium depletion.

To incorporate experimentally observed ROS-dependent ion channel remodeling into the Zukowski ECM-ROS model, I_NaL_ and I_CaL_ conductance scaling parameters were calibrated to experimental measurements obtained under oxidative stress conditions. Calibration of I_NaL_ and I_CaL_ was necessary to constrain ROS-dependent conductance scaling within physiologically observed ranges prior to whole-cell simulations.

Experimental measurements from Song et al. (2006) demonstrated 81.78 ± 24.98% increased late sodium current following exposure of guinea pig ventricular myocytes to 200µM H_2_O_2_ under a 300-ms voltage clamp protocol (Song *et al.,* 2006). To reproduce the experimental conditions *in silico*, cytosolic ROS concentration was increased to maximal simulated levels ([O ^.-^] ≥ 100µM), and I_NaL_ conductance (G_NaL_) scaling was adjusted such that simulated increases in I_NaL_ area under the curve (AUC) remained within the experimentally observed range at 1Hz (**Fig. 6A**). Representative steady-state traces at 1Hz demonstrated 103% AUC increase of I_NaL_ under high ROS conditions relative to no ROS conditions (**Fig. 6B**).

Similarly, ROS-dependent enhancement of I_CaL_ was calibrated using experimental measurements from Xie et al. (2009), in which adult rabbit ventricular myocytes exposed to 1mM H_2_O_2_ exhibited 65.75 ± 30.63% increased peak L-type calcium current during a 300-ms voltage clamp protocol. Under maximal simulated ROS conditions, I_CaL_ conductance (G_CaL_) scaling was tuned such that simulated peak I_CaL_ enhancement at 1Hz aligned with experimentally observed increases (**Fig. 6C**). Representative steady-state I_CaL_ traces at 1Hz demonstrated 45% increased peak calcium current during ROS exposure compared to no ROS conditions (**Fig. 6D**).

To validate model-predicted effects of ROS on electrophysiology and calcium handling, simulation predictions were compared to experimental measurements of hydrogen peroxide (H_2_O_2_) exposure in isolated ventricular myocytes. Guinea pig ventricular myocytes exposed to 200µM H_2_O_2_ with action potentials induced by a 5-ms depolarizing pulse at 0.16Hz exhibited APD_50_ prolongation of 37.25 ± 13.41% (Song *et al*., 2006) compared to no ROS. To assess model-predicted ROS effects across physiologically relevant pacing rates, cytosolic ROS ([O ^.-^] ) was held constant at 200µM and the model was paced for 50 beats at cycle lengths of 2000, 1000, 750, and 500ms (0.5-2Hz). Simulated APD_50_ was calculated as a percent increase from no ROS APD_50_. APD_50_ prolongation decreased with faster pacing, from 36.5% at 2000ms, 20.5% at 1000ms, 14.4% at 750ms, and 13.6% at 500ms cycle length (**Fig. 7A**). Song et al. (2006) paced myocytes at 0.16Hz (6250ms cycle length), far slower than the 0.5-2Hz (2000-500ms) range used in our simulations to reflect human heart rates from bradycardic to tachycardic. Simulated APD_50_ prolongation at 2000ms cycle length fell within the experimentally reported range (23.84-50.66%), while APD_50_ prolongation at 750ms, 1000ms, and 500ms fell below the experimental range which reflects the predicted rate-dependence of ROS-induced APD prolongation in the model.

**Figure 7:**
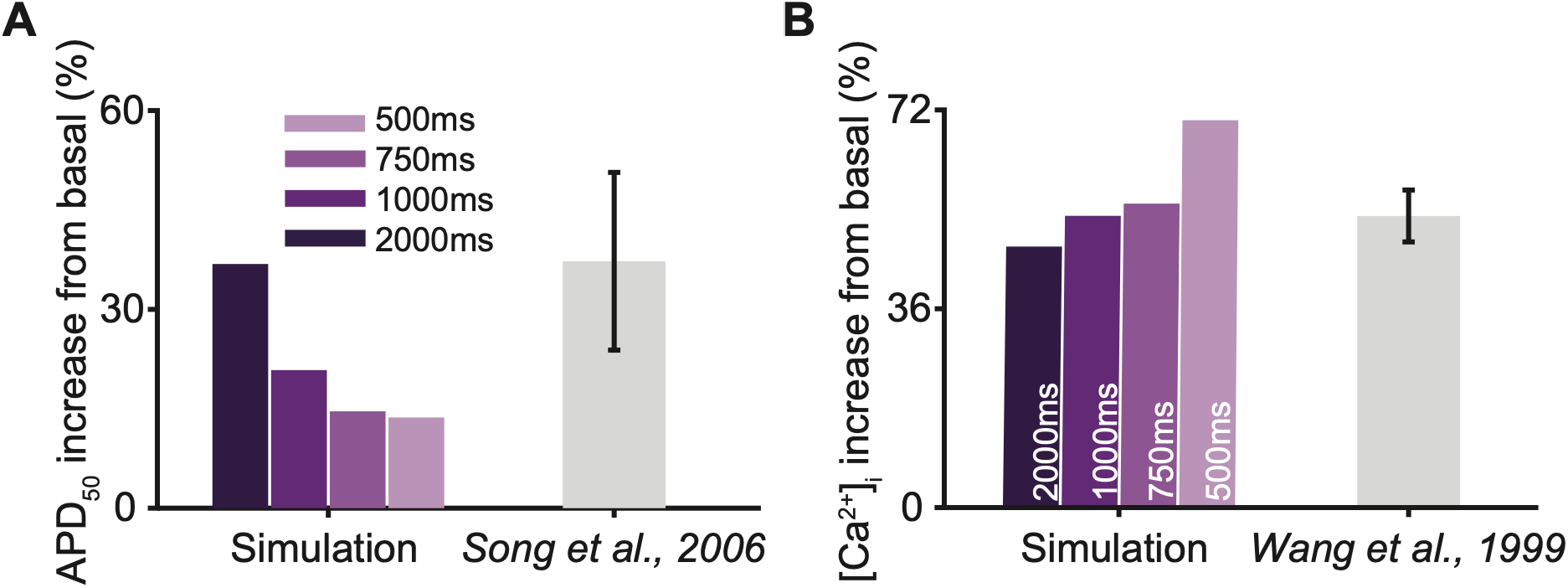
Validation of Zukowski ECM-ROS model-predicted ROS effects on action potential duration and cytosolic calcium. **(A)** Predicted percent increase in action potential duration at 50% repolarization (APD_50_) under high ROS conditions relative to baseline (no ROS) shown in colored bars. Experimental measurements (grey; Song *et al.,* 2006) were obtained from guinea pig ventricular myocytes exposed to 200µM H_2_O_2_ using a 5ms depolarizing pulse applied at 0.16Hz. Colored bars show simulated APD_50_ percent increases across pacing cycle lengths (CL = 2000, 1000, 750, 500ms). **(B)** Predicted percent increase in diastolic cytosolic calcium ([Ca^2+^]_i_) relative to baseline is shown in colored bars. Experimental measurements (grey; Wang *et al.,* 1999) were obtained from unpaced cell suspensions of Fura-2 loaded rat cardiomyocytes following incubation with 0.5mM H_2_O_2_. Error bars in experiments indicated standard deviation from the mean. Colored bars show simulated percent increases in [Ca^2+^]_i_ across pacing cycle lengths.

Model-predicted effects of ROS on calcium handling were validated against experimental measurements from Wang et al. (1999), in which isolated rat ventricular myocytes exposed to 0.5mM H_2_O_2_ exhibited a 52.8 ± 4.7% increase in cytosolic calcium AUC. To reproduce these conditions, cytosolic ROS was fixed at 0.5mM and the model was paced for 50 beats at cycle lengths of 2000, 1000, 750, and 500ms. Diastolic cytosolic calcium percent increase was calculated relative to the no ROS baseline at each pacing condition. Cytosolic calcium increased 46.6%, 52.0%, 54.5%, and 70.1% at cycle lengths of 2000ms, 1000ms, 750ms, and 500ms respectively (**Fig. 7B**). Simulated diastolic calcium percent increases across all pacing frequencies remained within or near the experimentally reported range of 52.8 ± 4.7%, except in the 2Hz condition which exceeded the upper experimental bound.

To investigate the integrated model-predicted effects of mitochondrial ROS production, the fraction of ROS that escaped the electron transport chain, which we refer to as “ETC ROS leak”, was varied from 0.40 to 0.60 across pacing cycle lengths (CL = 2000-500ms) and run to steady state. ETC ROS leak represents the fraction of electron transport chain flux (J_ETC_) contributing to mitochondrial ROS production ([O ^.-^] ). Modulation of ETC ROS leak alters steady-state ROS concentration, resulting in downstream effects on calcium handling through ROS-dependent modulation of the RyR, SERCA, I_CaL_, and I_NaL_ (**Fig. 8A**).

**Figure 8:**
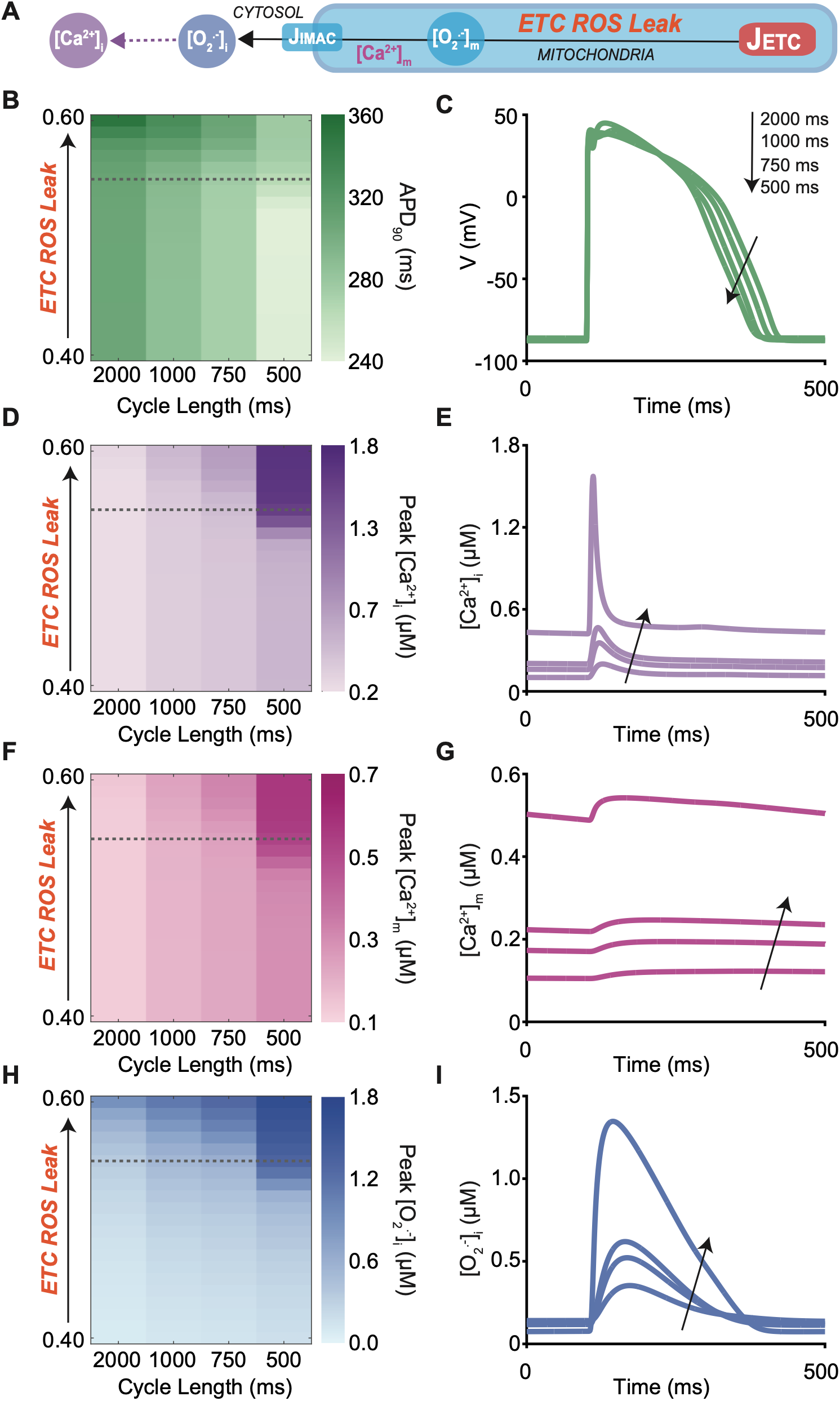
Predicted effects of ETC ROS leak and pacing on electrophysiology, calcium dynamics, and ROS dynamics in the Zukowski ECM-ROS model. **(A)** Schematic of mitochondrial ROS generation and transport. Electron transport chain (ETC) flux produces mitochondrial reactive oxygen species ([O_2_ ^.-^]_m_) as a fractional leak (ETC ROS leak). Mitochondrial [O_2_ ^.-^]_m_ is transported to the cytosol via the inner membrane anion channel (IMAC), contributing to cytosolic ROS ([O ^.-^]_i_). **(B, D, F, H)** Heat maps showing the steady-state dependence of APD_90_, peak cytosolic calcium ([Ca^2+^]_i_), peak mitochondrial calcium ([Ca^2+^]_m_), and peak cytosolic ROS ([O ^.-^] ), respectively, on ETC ROS leak (0.40-0.60, increment 0.01) and pacing cycle length. The dashed grey line at ETC ROS leak = 0.55 indicates the representative condition shown in time-domain traces. **(C, E, G, I)** Corresponding steady-state traces at ETC ROS leak = 0.55 across pacing cycle lengths (CL = 2000ms, 1000ms, 750ms, 500ms), with arrows indicating the direction of decreasing cycle length (toward 500ms). Across all variables, increasing ETC ROS leak and faster pacing interact to amplify electrophysiological and calcium dysregulation, with the most pronounced effects emerging at the combination of high leak and short cycle length.

Increasing ETC ROS leak produced progressive APD_90_ prolongation across all pacing frequencies (**Fig. 8B**). At CL=2000ms, APD_90_ increased from 305.0ms at ETC leak=0.40 to 356.0ms at ETC leak=0.60 (16.7% increase). At CL=500ms, APD_90_ increased from 241.6ms to 276.0ms (14.2% increase). Representative action potential traces at ETC leak=0.55 showed prediction of progressive APD prolongation with decreasing cycle length (**Fig. 8C**).

Peak cytosolic calcium predicted pronounced rate-dependent amplification with increasing ETC ROS leak (**Fig. 8D**). At CL=2000ms, peak [Ca^2+^]_i_ increased 41.3% from 0.188µM to 0.265µM across the ETC leak range. At CL=500ms, peak [Ca^2+^]_i_ increased 230.4% from 0.505µM to 1.669µM, which is more than five times greater than the slow pacing response. A marked nonlinear increase in peak [Ca^2+^]_i_ was observed at CL=500ms between ETC leak=0.40 (0.505µM) and ETC leak=0.55 (1.564µM), suggesting a pacing-dependent threshold for calcium overload at higher ROS levels. Representative cytosolic calcium transients at ETC leak=0.55 showed diastolic [Ca^2+^]_i_ increasing from 0.10µM at CL=2000ms to 0.42µM at CL=500ms (**Fig. 8E**).

Peak mitochondrial calcium similarly showed rate-dependent amplification, increasing 18.9% at CL=2000ms compared to 95.0% at CL=500ms across the ETC ROS leak range (**Fig. 8F**). Representative mitochondrial calcium transients at ETC ROS leak=0.55 showed diastolic [Ca^2+^]_m_ increasing from 0.11µM at CL=2000ms to 0.49µM at CL=500ms (**Fig. 8G**), consistent with progressive mitochondrial calcium accumulation at faster pacing rates driven by reduced diastolic recovery time.

Peak cytosolic ROS increased substantially across the ETC leak range, with peak [O_2_^.-^]_i_ increasing from 0.067µM to 1.000µM at CL=2000ms and from 0.165µM to 1.685µM at CL=500ms (**Fig. 8H**). Representative ROS traces at ETC ROS leak=0.55 showed peak [O_2_^.-^]_i_ increasing from 0.36µM at CL=2000ms to 1.35µM at CL=500ms, a 3.7-fold greater peak ROS response at faster pacing (**Fig. 8I**). Increased ETC ROS leak produced coordinated increases in APD prolongation, calcium accumulation, and ROS generation across all pacing conditions, with the largest absolute changes observed at faster pacing frequencies, consistent with rate-dependent amplification of the ROS-calcium positive feedback loop.

We next set out to characterize the predicted effects of doxorubicin (DOX) on mitochondrial and electrophysiological dysfunction across physiologically relevant pacing rates. Steady-state simulations were performed across DOX effect levels (D=0-1.0) at cycle lengths of 500, 750, and 1000ms. Consistent with the prediction of rate-dependent ROS amplification demonstrated in **Figure 8**, increasing DOX effect predicted progressive changes in all examined variables ([Ca^2+^]_i_, [Ca^2+^]_m_, ΔΨ, [O ^.-^] ). Faster pacing amplified the magnitude of predicted dysfunction and shifted the onset of nonlinear responses toward lower DOX effect levels.

Peak cytosolic calcium ([Ca^2+^]_i_) increased nonlinearly with DOX effect across all pacing rates **(Fig. 9A).** At 1000ms pacing, calcium elevation emerged above D=0.80, whereas at 500ms pacing, the onset shifted to ∼D=0.70. Peak mitochondrial calcium ([Ca^2+^]_m_) exhibited similar behavior, with greater concentrations at faster pacing rates and a nonlinear response that emerged at ∼D=0.70 **(Fig. 9B).** These results suggest that faster heart rates lower the DOX effect threshold at which calcium handling becomes compromised.

**Figure 9.**
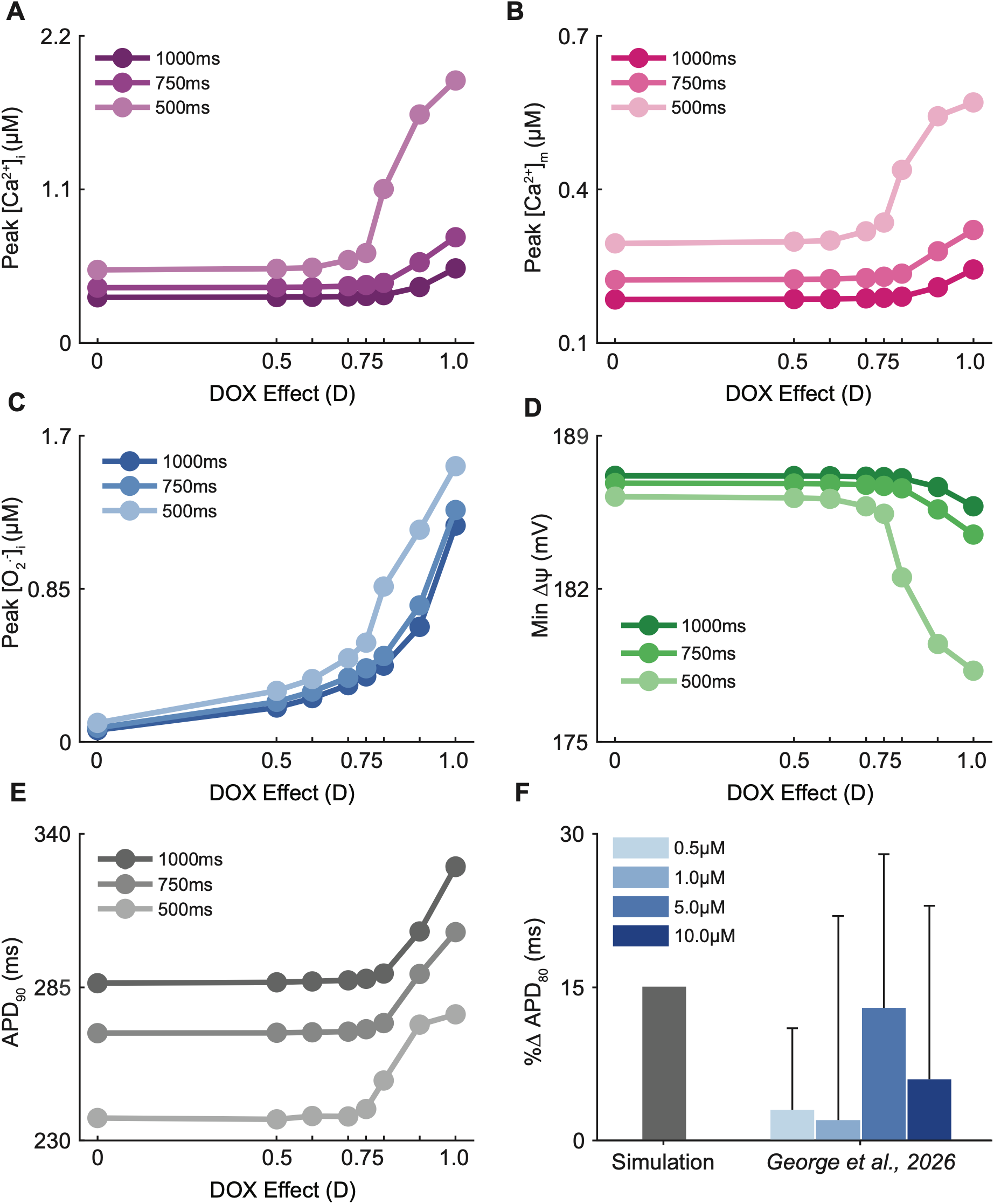
Predicted rate-dependent amplification of DOX-induced mitochondrial and electrophysiological dysfunction. Steady state model outputs plotted as a function of DOX effect (D=0–1.0) at cycle lengths of 1000ms (dark), 750ms (intermediate), and 500ms (light) for five variables: **(A)** peak cytosolic Ca^2+^ ([Ca^2+^]_i_, µM), **(B)** peak mitochondrial Ca^2+^ ([Ca^2+^]_m_, µM), **(C)** peak cytosolic superoxide ([O_2_^.-^]_i_, µM), **(D)** minimum mitochondrial membrane potential (ΔΨ, mV), **(E)** action potential duration at 90% repolarization (APD_90_, ms). All panels demonstrate faster pacing (500ms) amplifies DOX-induced dysfunction. **(F)** Validation of simulated APD_80_ (D=0 ◊ D=1, 1Hz) against experimental data from George et al. (2026), human ventricular slices from young male donors (n = 9, age = 39±9 years) treated with doxorubicin at increasing concentrations (0.5, 1, 5, and 10µM) for 24 hours. Error bars represent experimental +1 standard deviation.

Peak cytosolic ROS ([O_2_^.-^]_i_) demonstrated a comparable predicted pattern, with progressive elevation across DOX effect levels and the largest increases occurring at CL = 500ms **(Fig. 9C).** The rate-dependent shift in ROS threshold is consistent with the ROS-calcium positive feedback loop incorporated in the model, where reduced diastolic recovery time (inter pulse interval) at raster pacing rates promotes progressive calcium accumulation that drives further mitochondrial ROS production. Minimum mitochondrial membrane potential (ΔΨ) showed progressive depolarization with increasing DOX effect, again with 500ms pacing producing greater depolarization and an earlier threshold onset around D=0.6-0.7 at 500ms compared to D=0.75-80 at 1000ms **(Fig. 9D)**. Action potential duration at 90% repolarization (APD_90_) was predicted to exhibit rate-dependent threshold behavior consistent with the upstream mitochondrial and calcium perturbations **(Fig. 9E).** At 1000ms pacing, APD_90_ prolongation emerged above D=0.75-0.8, coinciding with the threshold for calcium and ROS elevation observed in panels A-D. At 500ms pacing, APD_90_ prolongation was predicted at D=0.6-0.7 consistent with rate-dependent ROS accumulation driving enhanced I_CaL_ and I_NaL_. Together, these results demonstrate that APD prolongation is a downstream consequence of DOX-induced mitochondrial dysfunction, amplified by faster pacing through the ROS-calcium positive feedback loop.

To validate the electrophysiological predictions for the Zukowski ECM-ROS model, simulated APD prolongation was compared with a human experimental dataset (**Fig. 9F**). Because the DOX effect parameter (D) represents a generalized level of DOX-induced stress rather than a specific pharmacological concentration, the maximal predicted effect (D=1.0) was compared with the full range of experimentally reported DOX concentrations, rather than a single matched dose. At 1Hz pacing, the model predicted 15.1% APD_80_ prolongation at full DOX effect (D=1.0). Human ventricular slices cultured with 0.5-10µM DOX for 24 hours exhibited APD_80_ prolongation of 2-13% across this concentration range in young male donor hearts (George et al., 2026). The maximal predicted effect was therefore comparable to the upper end of the experimentally observed response, which supports the physiological relevance of the electrophysiological predictions from the Zukowski ECM-ROS model.

To further investigate the temporal hierarchy between mitochondrial dysfunction and electrophysiological remodeling, simulations were performed at 2Hz steady-state pacing across increasing DOX effect levels (D=0.5-1.0) and percent changes from baseline (D=0) were quantified for each variable. While **Figure 9** demonstrated a common nonlinear threshold for APD prolongation and mitochondrial dysfunction, **Figure 10** shows substantial changes in mitochondrial variables at DOX effect levels where APD remains largely unaffected. This dissociation suggests mitochondrial dysfunction precedes detectable electrophysiological remodeling.

**Figure 10.**
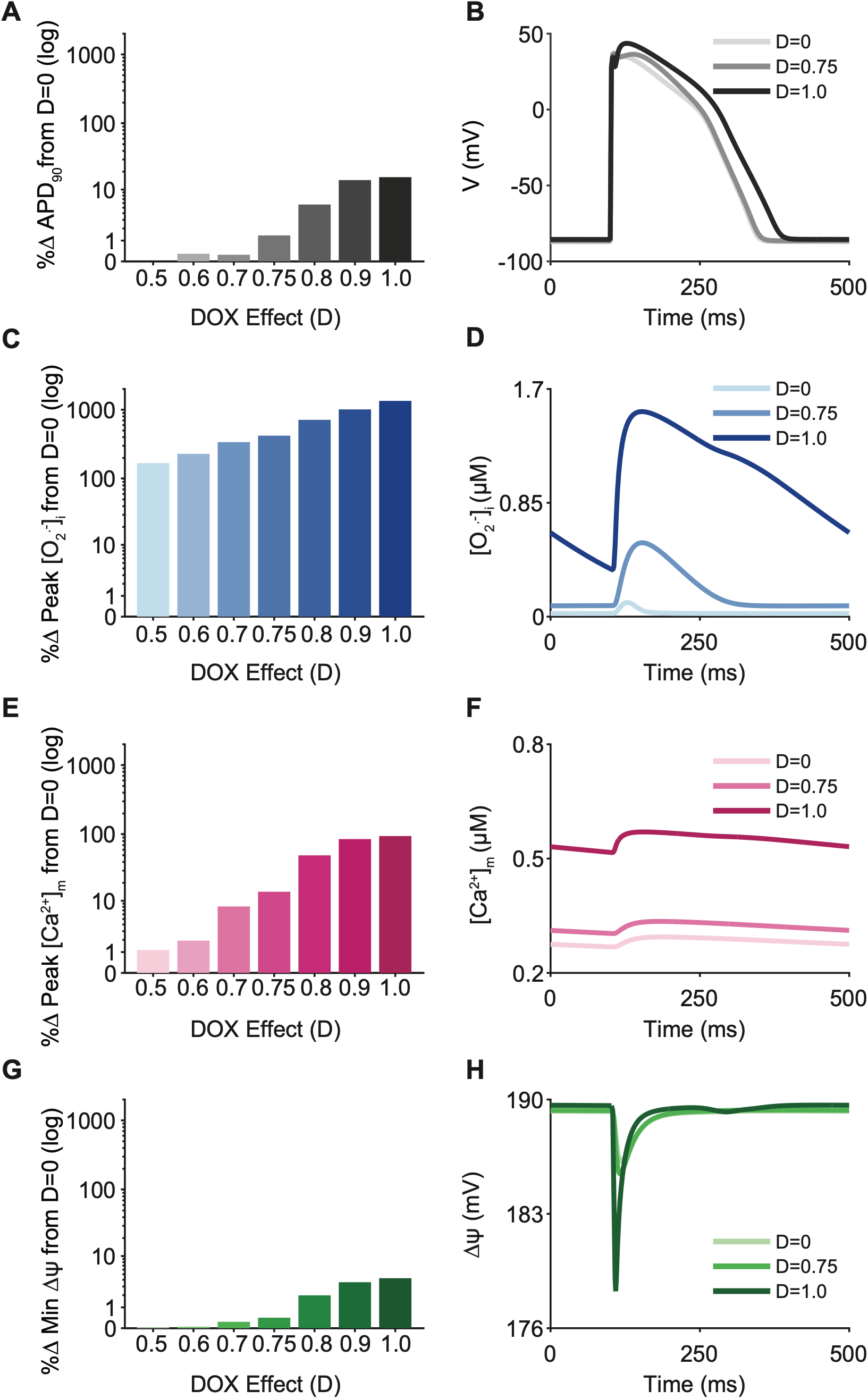
Mitochondrial dysfunction precedes electrophysiological remodeling across DOX effect levels. Simulations were performed at 2Hz pacing across increasing DOX effect (D=0.5-1.0). Left panels show percent change from no DOX (D=0) on a log scale. Right panels show representative steady state action potential and mitochondrial variable traces at D=0, D=0.75, and D=1.0. **(A, B)** APD showed minimal change at low-to-moderate DOX effect, with substantial prolongation only above D=0.75. **(C, D)** Cytosolic ROS ([O_2_ ^.-^]_i_) showed large increases beginning at D=0.5, with progressive accumulation across the action potential plateau. **(E, F)** Mitochondrial calcium ([Ca^2+^]_m_) showed early, graded elevation across DOX effect levels. **(G, H)** Mitochondrial membrane potential (ΔΨ) showed modest depolarization at high DOX effect.

APD showed minimal percent change across low-to-moderate DOX effect levels, increasing 0% at D=0.5 and 1.38% at D=0.75, with substantial prolongation emerging only above D=0.8 (5.67%) and reaching 15.6% at D=1.0 **(Fig. 10A, B)**. In contrast, cytosolic ROS ([O_2_ ^.-^]_i_) exhibited large percent increases beginning at D=0.5 (167%), with progressive accumulation reaching 338% at D=0.70, 420% at D=0.75, and 1343% at D=1.0 **(Fig. 10C, D).** Similarly, mitochondrial calcium ([Ca^2+^]_m_) exceeded the percent change in APD at every DOX effect level examined, though with substantially smaller magnitude than cytosolic ROS. Mitochondrial calcium increased 8% and D=0.70 and 14% at D=0.75 before a more pronounced nonlinear rise between D=0.8 (49%) and D=1.0 (94%), which reflects impaired mitochondrial calcium handling that precedes measurable electrophysiological change **(Fig. 10E, F)**. Mitochondrial membrane potential (ΔΨ) increased 0.03% at D=0.5 and reached a maximum depolarization of 4.3% at D=1.0 **(Fig. 10G, H)**. Mitochondrial membrane potential is tightly regulated, where even small depolarizations reflect mitochondrial dysfunction (Peoples *et al*., 2019; Detmer *et al*., 2022).

Together, these results demonstrate that mitochondrial ROS accumulation and calcium overload are potentially early indicators of DOX-induced cardiotoxicity, while action potential prolongation represents a later downstream consequence of sustained mitochondrial dysfunction. This temporal dissociation suggests that mitochondrial biomarkers may provide earlier warning of DOX-induced cardiotoxic stress than structural and functional cardiac changes, such as reduced ejection fraction and QT prolongation (Lotrionte *et al*., 2013; Benjanuwattra *et al*., 2020).

To investigate how inter-individual variability in ion channel expression modulates arrhythmia vulnerability under DOX stress, a population of N=1000 virtual human ventricular cells with ±20% randomized variability across 12 ion channel conductance scaling factors was subjected to a pause protocol. Cells were paced for 10 beats at 2Hz followed by a 1 second pause, and the post-pause beat was analyzed.

Common medications co-administered with DOX in cancer patients have been shown to prolong the QT interval through block of the rapid delayed rectifier potassium current (I_Kr_) (Agnihotri *et al*., 2024; Ramasubbu *et al*., 2025). To examine the progression of I_Kr_ block effects in the context of DOX-induced oxidative stress, simulations were performed across five conditions: no DOX with 0% I_Kr_ block, full DOX (D=1) with 0%, 30%, and 40% I_Kr_ block, and no DOX with 40% I_Kr_ block.

Under baseline conditions (D=0, 0% I_Kr_ block), the population exhibited tight clustering of action potential morphology and values, as well as mitochondrial variables, with mean APD_90_ of 253.3 ± 11.3ms **(Fig. 11A, F, K, P)**. I_Kr_ block alone (D=0, 40% I_Kr_ block) was predicted to prolong mean APD_90_ to 332.5 ± 15.6ms. Notably, all cells repolarized normally with no EADs, and mitochondrial calcium and cytosolic ROS remained near baseline levels **(Fig. 11E, J, O, T)**. DOX alone (D=1, 0% I_Kr_ block) prolonged mean APD_90_ to 255.6 ± 10.4ms with elevated mitochondrial calcium and cytosolic ROS, yet all cells also repolarized normally with no EADs **(Fig. 11B, G, L, Q)**. These results demonstrate the model prediction that neither DOX-induced oxidative stress nor I_Kr_ block alone is sufficient to produce EADs.

**Figure 11:**
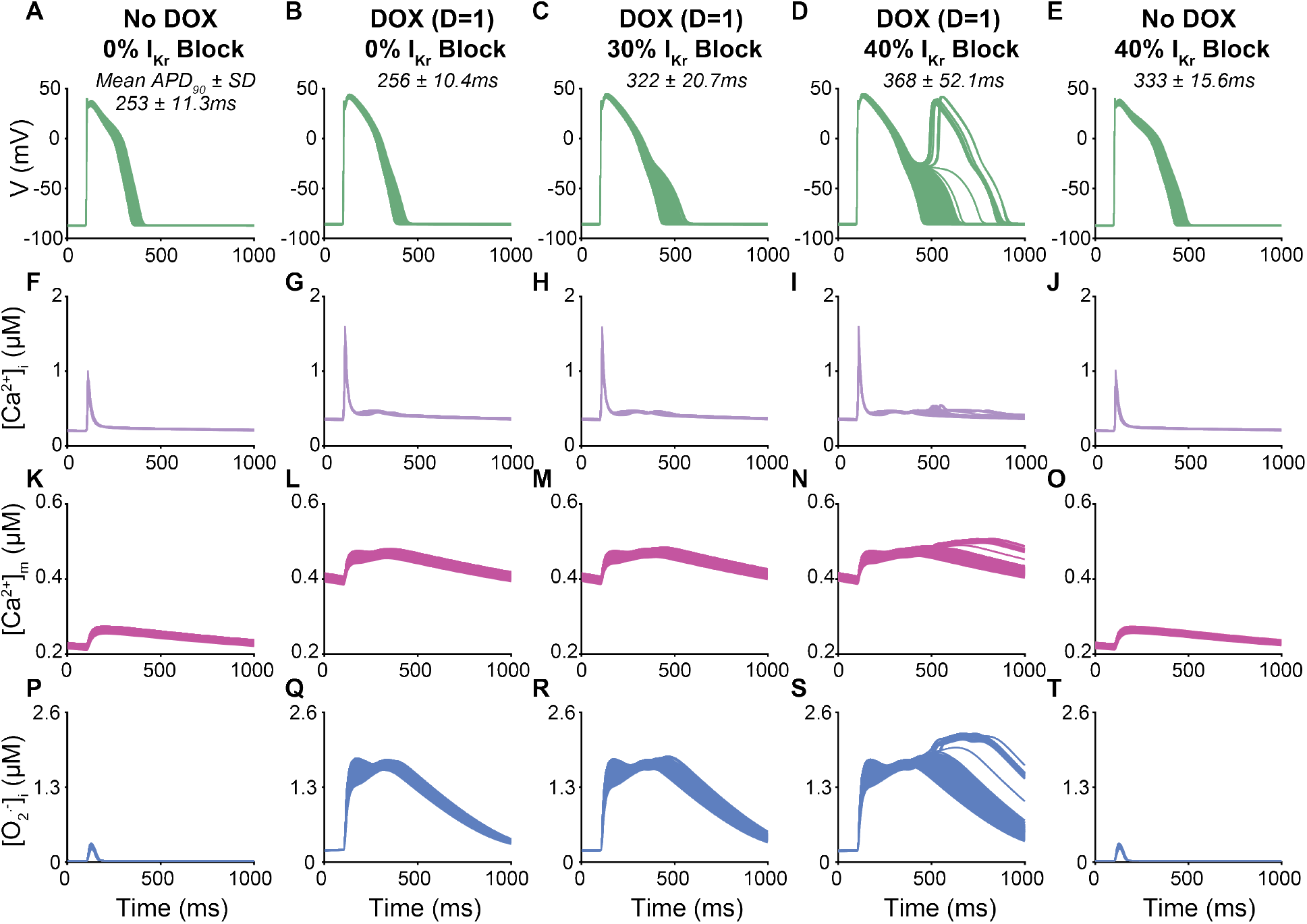
Predicted 2Hz population traces across DOX and I_Kr_ block conditions. A population of N=1000 virtual human ventricular cells with ±20% random variability across 12 ion channel conductance scaling factors was paced at 2Hz for 10 beats followed by a 1 second pause. The post-pause beat is shown. Mean APD_90_ ± standard deviation (SD) is annotated on each voltage panel. Each column shows membrane potential (V, mV), cytosolic calcium ([Ca^2+^]_i_, µM), mitochondrial calcium ([Ca^2+^]_m_, µM), and cytosolic ROS ([O_2_^.-^]_i_, µM) from top to bottom. No DOX, 0% I_Kr_ block: **(A, F, K, P)**. Full DOX effect (D=1), 0% I_Kr_ block: **(B, G, L, Q)**. Full DOX effect (D=1), 30% I_Kr_ block: **(C, H, M, R)**. Full DOX effect (D=1), 40% I_Kr_ block: **(D, I, N, S)**. No DOX, 40% I_Kr_ block: **(E, J, O, T)**.

Under combined DOX and I_Kr_ block, progressive I_Kr_ block further prolonged APD_90_ and increased population variability. At 30% I_Kr_ block, mean APD_90_ increased to 322.1 ± 20.7ms with a broad spread across the population, though all cells continued to repolarize normally **(Fig. 11C, H, M, R)**. At 40% I_Kr_ block, mean APD_90_ increased to 367.5 ± 52.1ms and a subset of the population developed EADs, visible as secondary voltage upstrokes during the repolarization phase **(Fig. 11D, I, N, S)**. The large SD at this condition reflects the heterogeneous population response where some cells repolarized normally while others exhibited prolonged or uncompleted repolarization. Notably, despite comparable APD_90_ between the no DOX 40% I_Kr_ block condition (332.5 ± 15.6ms) and the DOX 30% I_Kr_ block condition (322.1 ± 20.7ms), EADs were absent without DOX, confirming that APD prolongation alone is insufficient and that DOX-induced oxidative stress is a necessary condition for EAD emergence.

## DISCUSSION

The Zukowski ECM-ROS model is, to our knowledge, the first computational model to couple the O’Hara-Rudy (ORd) human ventricular electrophysiological framework to mitochondrial energetics and ROS dynamics (**Fig 1)**. The model includes a ROS-Ca^2+^ positive feedback loop in which cytosolic calcium enters the mitochondria and drives superoxide production (ROS) as a fraction of electron transport chain flux. Cytosolic ROS modulates the ryanodine receptor, SERCA, L-type calcium current, and late sodium current and thereby alters calcium handling and action potential duration. The model reproduces experimental measurements of mitochondrial calcium uptake and ROS-dependent changes in calcium and action potential duration.

We used the Zukowski ECM-ROS model to investigate the early mechanisms of doxorubicin (DOX)-induced cardiotoxicity. Simulations suggest that mitochondrial dysfunction and ROS-dependent calcium and ion channel modulation are sufficient to produce DOX-induced action potential prolongation. Moreover, the model predicts that acute application of DOX leads to rate-dependent amplification of mitochondrial and electrophysiological dysfunction, as well as temporal dissociation where mitochondrial ROS accumulation precedes measurable action potential prolongation. We also assessed the effects of DOX-induced oxidative stress within a variable cell population to mimic physiological variability in human cardiac myocytes (Muszkiewicz *et al*., 2016; Trayanova *et al*., 2024). Because DOX is often administered with drugs that reduce repolarization reserve, like ondansetron (Zofran), we tested the effects DOX and I_Kr_ block alone and in combination. The simulations suggest that DOX alone and low-level drug block of I_Kr_ is individually insufficient, but jointly sufficient, to produce early afterdepolarizations, consistent with clinical data suggesting electrical disruption in some patients (Kilickap *et al*., 2007; Doggrell & Hancox, 2013; Freedman *et al*., 2014).

Simulations in the Zukowski ECM-ROS model suggest that mitochondrial calcium transients closely track with cytosolic calcium dynamics, and steady-state mitochondrial calcium uptake match experimental measurements across a physiological range of cytosolic calcium concentrations (**Fig. 2**). Together, these data validate and support plausibility of the model formulation of calcium exchange between the cytosol and mitochondrial matrix. Validation of the mitochondrial module was performed on the calcium handling data available in the literature at the time of model development, which is derived from rat ventricular myocytes (Andrienko *et al*., 2009) and human HeLa cells, a human cell line derived from cervical cancer cells taken from Henrietta Lacks (Collins *et al*., 2001) (**Fig. 2B**). Mitochondrial and ROS module parameter values (**Table 1**) were adapted directly from prior computational models (Cortassa *et al*., 2003, 2004; Zhou *et al*., 2009; Senneff & Lowery, 2022). Mitochondrial bioenergetic pathways, including the TCA cycle and oxidative phosphorylation complexes, are broadly conserved across mammalian cells, which supports the general applicability of the rate law structures adapted in the Zukowski ECM-ROS model (Arnold & Finley, 2023).

The steady-state mitochondrial fluxes in the Zukowski ECM-ROS model describe calcium uptake and efflux, metabolic activity, electron transport chain flux, ATP production and transport, and proton leak without a full biochemical representation of mitochondrial metabolism (**Fig. 3**). More detailed mitochondrial models include individual TCA cycle intermediates, oxidative phosphorylation pathways, redox states, and, in some cases, spatial organization of mitochondrial and cellular compartments (Aon *et al*., 2003; Zhou *et al*., 2009; Li *et al*., 2015). This level of detail is necessary when the biological question depends on a specific metabolic pathway or mitochondrial process. For example, detailed models can distinguish the contributions of individual respiratory complexes, identify changes in metabolic substrates or redox state, resolve ATP depletion during severe energetic stress, or examine spatial gradients in ROS and calcium between mitochondria and the sarcoplasmic reticulum. These models are therefore well suited to questions that require identification of the specific biochemical or spatial origin of mitochondrial dysfunction.

The Zukowski ECM-ROS model addresses a different question. The objective of the present study is to determine how changes in mitochondrial ROS production affect calcium handling and human ventricular electrophysiology at the whole-cell level. The model therefore represents mitochondria as a single, well-mixed compartment and uses a reduced set of fluxes to describe the major pathways that connect mitochondrial calcium, electron transport chain activity, ROS production, and cellular electrophysiology. This structure allows individual processes, such as ETC ROS leak or cytosolic calcium, to be perturbed directly and their effects on ROS accumulation, calcium overload, and action potential duration to be isolated. The reduced formulation also limits computational cost, which enables population-scale simulations. Additional biochemical or spatial detail can be incorporated in future versions when required to address a different biological question or context of use.

A central prediction of this study is that mitochondrial ROS production is sensitive to changes in cytosolic calcium and that this relationship is strongly modulated by pacing rate. A 62% increase in peak cytosolic calcium produced a 4.3% increase in peak mitochondrial calcium, an 18.8% increase in electron transport chain flux, and a 52.6% increase in cytosolic ROS (**Fig. 4B-E**). Thus, relatively modest increases in mitochondrial calcium and electron transport chain activity produced disproportionately large increases in ROS. This nonlinear response is consistent with the role of ROS as a sensitive physiological signal rather than solely a marker of severe oxidative stress (Sies & Jones, 2020). The nonlinear sensitivity to cytosolic calcium was further modulated by pacing rate. The same 50% cytosolic calcium perturbation produced a 2.66-fold greater ROS response at 500ms compared to 2000ms cycle length, and this ratio remained consistent across all tested calcium levels (**Fig. 4F**). Together, these findings reveal a rate dependent coupling between cytosolic calcium loading and mitochondrial ROS production. The model predicts that pacing frequency acts as a critical modifier of mitochondrial vulnerability and amplifies the oxidative consequences of altered calcium handling. This relationship provides a mechanistic framework for how relatively modest disturbances in cellular calcium homeostasis may produce substantial oxidative stress under conditions of increased cardiac demand.

A second key finding is that the effects of ROS on SR calcium handling are biphasic rather than uniformly detrimental. The model predicted that at moderate ROS concentrations, ROS-dependent RyR activation increased calcium release and cytosolic calcium transient amplitude. At higher ROS concentrations, calcium stores in both the NSR and JSR were depleted, resulting in reduced calcium cycling (**Fig. 5**). This concentration-dependent response parallels the biphasic pattern reported by Sag et al. (2013) following cardiac irradiation. Acute irradiation increased calcium transient amplitude and SR calcium reuptake, whereas chronic irradiation one week later reduced calcium transient amplitude and SR calcium load (Sag *et al*., 2013). Sag et al. (2013) attributed the temporal biphasic effect to ROS-dependent activation of CaMKII and subsequent phosphorylation of RyR2 and phospholamban (Sag *et al*., 2013). The Zukowski ECM-ROS model does not explicitly represent CaMKII activation kinetics. Instead, ROS-dependent modulation of RyR and SERCA was calibrated to reproduce net experimental effects associated with oxidative stress. The resulting biphasic response is consistent with the distinction between physiological redox signaling and pathological oxidative stress (Sies & Jones, 2020; Zukowski & Clancy, 2026). Moderate ROS elevation may act as an adaptive signal and transiently enhance calcium release, while sustained or substantial ROS elevation overwhelms the compensatory response and drives SR calcium store depletion.

Beyond its effects on SR calcium handling, oxidative stress also alters membrane currents that shape the action potential and intracellular calcium loading. We therefore incorporated ROS dependent enhancement of I_CaL_ and I_NaL_ to capture the broader electrophysiological consequences of increased oxidative stress (**Fig. 6**). ROS-dependent scaling functions for I_CaL_ and I_NaL_ were fit to net experimental effects, rather than modeling intermediate signaling steps explicitly (**Fig. 6**). In the model, cytosolic superoxide modulates RyR, SERCA, I_CaL_, and I_NaL_. However, the effects on these four targets were calibrated using experimental data from exogenous H_2_O_2_ exposure. In the absence of equivalent superoxide specific data, experimental responses to exogenous H_2_O_2_ were used as a proxy to calibrate the effects of cytosolic oxidative stress on RyR, SERCA, I_CaL_ and I_NaL_ through scaling functions (Ransy *et al*., 2020).

Experimental data indicate that ROS exposure increases peak I_CaL_ and slows current inactivation (Xie *et al*., 2009; Song *et al*., 2011). In the model, increasing I_CaL_ permeability reproduces the increase in peak current but only modestly slows inactivation. A larger permeability increase would better reproduce the observed slowing of inactivation but would overestimate the peak current. Thus, permeability scaling alone cannot simultaneously capture both experimentally observed effects. Reproducing both the increased peak and slowed inactivation observed experimentally will require future modification of the kinetics of the current directly, for example through ROS-dependent changes in activation or inactivation gating variables.

To determine whether the integrated model reproduces experimentally observed effects of ROS on cardiac electrophysiology and calcium handling, we compared its predictions with independent experimental data (**Fig. 7**). Simulated APD_50_ prolongation fell within the experimental range (Song *et al*., 2006) at 0.5Hz pacing. The available experimental data were collected in guinea pig myocytes at 0.16Hz, which is substantially slower than the 0.5-2Hz range simulated to reflect human heart rates (**Fig. 7A**). Simulated cytosolic calcium accumulation closely matched experimental measurements (Wang *et al*., 1999) across nearly the full range of pacing rates tested, with the fastest rate (2Hz) exceeding the upper bound of the experimentally reported range (**Fig. 7B**). The experimental data from Wang et al. (1999) were obtained from unpaced rat ventricular myocytes at room temperature, whereas the model simulated diastolic calcium in human ventricular myocytes during continuous pacing. Differences in species, temperature, and pacing conditions likely explain the deviations at faster rates. Further validation will require direct measurements of ROS dependent APD prolongation and calcium accumulation in human cardiac tissue across physiologically relevant pacing rates.

Having established that the model reproduced experimentally observed effects of ROS on action potential duration and calcium accumulation, we next simulated how mitochondrial ROS production and pacing rate interact to drive electrophysiological and calcium dysregulation independent of a specific pathological trigger (**Fig. 8**). We did this by varying ETC ROS leak across a range of pacing cycle lengths and quantifying the resulting changes in cytosolic ROS, calcium handling, and action potential duration. Zhou et al. (2009) used a similar approach, varying a ROS “shunt” parameter to characterize a model response to oxidative stress, but predicted APD shortening rather than prolongation. The contrasting predictions of APD shortening by Zhou et al. (2009) and APD prolongation by the Zukowski ECM ROS model reflect fundamental differences in the mechanisms represented by each model. Zhou et al. (2009) predicted increased ROS production drives mitochondrial membrane potential collapse, which increases the cytosolic ADP/ATP ratio, activates the sarcolemmal ATP-sensitive potassium current, and shortens the APD. This pathway is relevant to severe, acute energetic collapse such as ischemia-reperfusion (Akar *et al*., 2005; Zhou *et al*., 2009).

Here, our intent was to capture the ROS-calcium feedback loop in which elevated ROS increases RyR calcium release, decreases SERCA calcium uptake, increases I_CaL_ and I_NaL_, and results in increased cytosolic calcium and APD prolongation. The Zukowski ECM-ROS model does not include cytosolic ATP/ADP dynamics or a K_ATP_ current, so ROS-driven APD change arises entirely through direct modulation of I_CaL_, I_NaL_, RyR, and SERCA. The Zukowski ECM-ROS model is well suited as a generalizable framework for studying ROS-driven calcium and electrophysiological dysfunction prior to energetic collapse across a range of conditions. Additional pathways, such as ATP/ADP dynamics or metabolically sensitive currents, can be incorporated in future model instances to extend model applicability in new domains.

Previously published computational models of DOX-induced cardiotoxicity have incorporated drug effects through direct modification of ion channel conductance (Fernandez-Chas *et al*., 2018). Fernandez-Chas et al. (2018) derived consensus scaling factors from published experimental data and applied them directly to the I_Kr_, I_CaL_, sarcoplasmic reticulum calcium leak current, I_NaK_, I_NaCa_, SERCA, and RyR as functions of DOX or doxorubicinol concentration. This approach reproduced the effects of acute and chronic DOX exposure on action potential duration and calcium handling in rabbit and human ventricular models. However, because it represents the net effects of DOX on each target, it cannot distinguish direct drug interactions from changes arising through upstream mechanisms or predict how electrophysiological remodeling evolves with the severity of mitochondrial oxidative stress. The Zukowski ECM-ROS model addresses this limitation by representing DOX effects through increased mitochondrial ROS production and reduced antioxidant scavenging, thereby linking oxidative stress mechanistically to the resulting electrophysiological phenotype.

We next evaluated whether this mechanistic representation of DOX induced oxidative stress could reproduce the experimentally observed electrophysiological response to DOX exposure. Simulated APD_80_ prolongation at maximal DOX effect fell within the range of experimentally reported values across DOX concentrations from 0.5-10µM (**Fig. 9F**, George *et al.,* 2026). The DOX effect parameter (D) captures a generalized level of ETC-dependent ROS production and antioxidant depletion, but neither of these underlying mechanistic parameters have been experimentally linked to a specific micromolar DOX concentration in cardiomyocytes. As a result, the model cannot currently predict which concentration of DOX corresponds to any given value of D. Nonetheless, the simulated maximal DOX effect (D=1) predicted APD_80_ prolongation within the experimental range, which provides confidence in the simulated DOX-induced electrophysiological predictions.

The model was applied to simulate acute DOX-induced oxidative stress across a range of pacing rates. The degree of DOX-induced oxidative stress necessary to produce notable rate-dependent calcium, ROS, and APD effects changed from D≈0.75-0.80 at 1000ms pacing to D≈0.60-0.70 at 500ms pacing (**Fig. 9A-E**). The pacing shift has direct clinical relevance since patients undergoing chemotherapy frequently experience tachycardia independently of, or concurrently with, DOX-induced cardiotoxicity (Sakellakis *et al*., 2024; Fakih *et al*., 2025). The model predictions suggest that physiological or clinical tachycardia may lower the threshold DOX exposure at which cardiotoxic dysfunction becomes detectable, independent of any change in cumulative DOX dose.

The temporal relationship between mitochondrial and electrophysiological dysfunction was predicted to be dissociated. At a DOX effect level (D=0.5), APD was unchanged, whereas cytosolic ROS increased by 167%, and mitochondrial calcium increased by 1%. Substantial APD prolongation did not emerge until D≥0.8 (**Fig. 10**). This suggests that ROS accumulation precedes detectable electrophysiological remodeling during the progression of DOX-induced mitochondrial dysfunction. These results identify increased ROS as a potential early indicator of DOX cardiotoxicity before changes in action potential duration become detectable. Current clinical monitoring for DOX cardiotoxicity relies on downstream structural and functional cardiac markers, including reduced ejection fraction and QT interval prolongation (Plana *et al*., 2014).

The model predicts that oxidative stress may develop before these conventional markers of cardiac dysfunction become detectable. This finding is consistent with growing clinical interest in circulating biomarkers of oxidative stress and mitochondrial injury as earlier indicators of cardiotoxicity (Murtagh *et al*., 2023; Kim *et al*., 2025; Kong *et al*., 2026). Future experimental and clinical studies are needed to determine whether specific concentrations of ROS or mitochondrial calcium can identify early DOX-induced cardiac dysfunction and to define clinically relevant thresholds.

We next tested whether DOX-induced oxidative stress increases arrhythmia susceptibility in the presence of reduced repolarization reserve. This interaction has direct clinical relevance because ondansetron (Zofran), a serotonin 5-HT3 receptor antagonist commonly administered as an antiemetic during DOX treatment, also blocks I_Kr_ through inhibition of the hERG channel (Keefe, 2002; Georgy *et al*., 2007; Barni *et al*., 2016; Singh *et al*., 2023). Because acute proarrhythmia in patients with combined DOX treatment and I_Kr_ block is rare, we developed a population of models with electrophysiological parameters varied within experimentally reported ranges to identify conditions that promote arrhythmia. Population model predictions suggest that DOX-induced oxidative stress and pharmacological I_Kr_ block alone prolong mean APD_90_ to 255.6ms and 332.5ms, respectively, compared with 253.3ms at baseline. We tested 0%, 30%, and 40% I_Kr_ block conditions in the population model to represent a range of clinically plausible reductions in repolarization reserve (**Fig. 11**). However, key arrhythmia markers, namely early afterdepolarizations (EADs), emerged only with combined DOX and 40% I_Kr_ block. These results suggest that DOX-induced oxidative stress increases susceptibility to EADs when repolarization reserve is reduced and that this risk may not be fully captured by QT interval monitoring alone. This prediction is consistent with postmarketing surveillance data showing that reported arrhythmias temporally associated with ondansetron administration occur predominantly in patients with a concomitant QT-prolonging medication or significant cardiac history (Kilickap *et al*., 2007; Doggrell & Hancox, 2013; Freedman *et al*., 2014; Tafish *et al*., 2025).

### Limitations

Several limitations should be considered when interpreting the predictions from the Zukowski ECM-ROS model. First, parameters within the mitochondrial and ROS modules were adapted from prior computational models and validated against the experimental data available at the time of model development. Some of these data were obtained from non-human or non-cardiac cells. Therefore, the extent to which the specific kinetic parameters quantitatively represent human cardiac mitochondria has not been directly established. Validation against human cardiac mitochondrial data represents an important target for future model refinement.

Second, the present model focuses on mitochondrial ROS-dependent mechanisms of DOX cardiotoxicity and does not include direct effects of DOX on repolarizing potassium currents. Experimental measurements in a heterologous expression system indicate that DOX inhibits I_Ks_ at low micromolar concentrations, with limited direct effect on I_Kr_ at concentrations up to 30µM (Ducroq *et al*., 2010). We excluded direct I_Ks_ inhibition to isolate the contribution of mitochondrial dysfunction and ROS-dependent calcium and ion channel modulation to DOX-induced electrophysiological remodeling. The model reproduced experimental APD prolongation through ROS-dependent mechanisms alone, but direct DOX inhibition of I_Ks_ could further reduce repolarization reserve and increase arrhythmia susceptibility. Future model extensions can incorporate a direct, dose-dependent effect of DOX on I_Ks_ as additional experimental data become available.

Third, ROS-dependent modulation of RyR, SERCA, I_CaL_, and I_NaL_ was calibrated from experiments that applied extracellular H_2_O_2_, whereas cytosolic superoxide is the ROS that directly modifies these targets in the model. Extracellular H_2_O_2_ concentration cannot be quantitatively mapped to intracellular superoxide concentration. These relationships therefore represent the net effects of oxidative stress rather than a direct correspondence between the concentrations of H_2_O_2_ and O_2_^.-^. In addition, the I_CaL_ and I_NaL_ scaling functions were constrained by single maximal-effect measurements because full concentration-response data are not currently available. ROS-dependent effects on I_CaL_ and I_NaL_ were represented through changes in permeability and conductance, respectively, rather than direct modification of channel kinetics. Additional concentration-dependent and species-specific experimental measurements will allow these relationships to be refined.

The current limitations primarily affect quantitative interpretation of specific parameter values and thresholds rather than the intended use of the model as a reduced framework to investigate interactions between mitochondrial ROS, calcium handling, and human ventricular electrophysiology. Future experimental data can be incorporated to refine individual mechanisms while retaining the integrated structure of the model.

## Conclusions

This study presents the first computational model coupling human ventricular electrophysiology to mitochondrial energetics and ROS dynamics, and the first to reproduce DOX-induced electrophysiological remodeling resulting from acute oxidative stress. Beyond model development, key model predictions include, 1) faster pacing amplifies mitochondrial and electrophysiological dysfunction, 2) ROS accumulation precedes action potential prolongation, and 3) DOX increases susceptibility to arrhythmia in the presence of I_Kr_ block. These findings offer testable hypotheses with direct relevance to the timing of cardiotoxicity surveillance and the selection of co-administered medications in oncology practice. Model predictions emerged from a simplified, single-compartment mitochondrial and ROS description that retains the components necessary and sufficient to capture the ROS-calcium feedback loop. The necessary but sufficient complexity in the model design constitutes an extensible framework for future inclusion of ATP/ADP dynamics and additional ROS-sensitive pathways to address specific pathological questions.

## Author Contributions

H.M.Z. developed and implemented the computational model, conducted the simulations, analyzed and visualized the results, and drafted the manuscript. P.C.Y. contributed to model development, simulation design, and interpretation of the results. G.H.H. contributed experimental and physiological expertise and interpretation of the results. C.E.C. conceived and supervised the study, contributed to model design and interpretation of the results, and revised the manuscript. All authors reviewed and approved the final manuscript.

## Generative AI Statement

For the manuscript, the authors used ChatGPT (OpenAI, GPT-5.6) and Claude (Anthropic, Sonnet 5) for language editing purposes. These tools were not used for model development, generation of results, or scientific interpretation. All AI-assisted text was reviewed and edited by the authors, who take full responsibility for the content of the manuscript.

## Funding

Dr. Colleen E. Clancy is supported in part by the National Institutes of Health under grants R01HL128537, OT2OD026580, and R01HL17400 (to C. E. Clancy). Additional support was provided to C. E. Clancy by The UC Davis Center for Precision Medicine and Data Sciences and its computing facilities. Hannah M. Zukowski was supported by the National Institutes of Health grant T32HL086450 (to H. M. Zukowski).

## Notes

### Competing Interest Statement

The authors have declared no competing interest.

